# *TSC2* loss in neural progenitor cells suppresses translation of ASD/NDD-associated transcripts in an mTORC1- and MNK1/2-reversible fashion

**DOI:** 10.1101/2024.06.04.597393

**Authors:** Pauline Martin, Krzysztof J. Szkop, Francis Robert, Srirupa Bhattacharyya, Roberta L. Beauchamp, Jacob Brenner, Nicholas E. Redmond, Sidong Huang, Serkan Erdin, Ola Larsson, Vijaya Ramesh

## Abstract

Tuberous sclerosis complex (TSC) is an inherited neurodevelopmental disorder (NDD) with frequent manifestations of epilepsy and autism spectrum disorder (ASD). TSC is caused by inactivating mutations in *TSC1* or *TSC2* tumor suppressor genes, with encoded proteins hamartin (TSC1) and tuberin (TSC2) forming a functional complex inhibiting mechanistic target of rapamycin complex 1 (mTORC1) signaling. This has led to treatment with allosteric mTORC1 inhibitor rapamycin analogs (“rapalogs”) for TSC tumors; however, rapalogs are ineffective for treating neurodevelopmental manifestations. mTORC1 signaling controls protein synthesis by regulating formation of the eIF4F complex, with further modulation by MNK1/2 kinases via phosphorylation of the eIF4F subunit eIF4E. While both these pathways modulate translation, comparing their impact on transcriptome-wide mRNA translation, as well as effects of inhibiting these pathways in TSC has not been explored. Here, employing CRISPR-modified, isogenic TSC2 patient-derived neural progenitor cells (NPCs), we have examined transcriptome-wide changes in mRNA translation upon *TSC2* loss. Our results reveal dysregulated translation in *TSC2*-Null NPCs, which significantly overlaps with the translatome from *TSC1*-Null NPCs. Interestingly, numerous non-monogenic ASD-, NDD-and epilepsy-associated genes identified in patients harboring putative loss-of-function mutations, were translationally suppressed in *TSC2*-Null NPCs. Importantly, translation of these ASD- and NDD-associated genes was reversed upon inhibition of either mTORC1 or MNK1/2 signaling using RMC-6272 or eFT-508, respectively. This study establishes the importance of mTORC1-eIF4F- and MNK-eIF4E-sensitive mRNA translation in TSC, ASD and other neurodevelopmental disorders laying the groundwork for evaluating drugs in clinical development that target these pathways as a treatment strategy for these disorders.

## INTRODUCTION

Tuberous sclerosis complex (TSC [MIM: 191100 and MIM: 613254]) is an autosomal dominant neurodevelopmental disorder (NDD) with clinical involvement of multiple organ systems including the central nervous system (CNS). TSC is caused by mutations in the *TSC1* [MIM: 605284] or *TSC2* [MIM: 191092] gene, encoding the tumor suppressor protein hamartin (TSC1) or tuberin (TSC2), respectively. In addition to slow growing hamartomas in many organs, neurodevelopmental manifestations such as epilepsy, intellectual disability and autism spectrum disorders (ASD) are commonly observed in these patients ^1–3^. TSC proteins function as a complex, relaying signals from diverse cellular pathways to control mechanistic target of rapamycin (mTOR [MIM: 601231]) complex 1 (mTORC1) signaling. Loss of TSC1/TSC2 proteins results in aberrant activation of mTORC1, which has led to first-generation mTORC1 rapalog inhibitors, emerging as a lifelong therapy for TSC hamartomas ^4–7^. Moreover, recent clinical trials reveal that rapalogs reduce epilepsy in 40% of TSC patients ^8^. In contrast, rapalogs are ineffective in treating TSC-associated neuropsychiatric disorders (TAND) and autism ^9; 10^.

Among its many activities, mTORC1 signaling plays an essential role in protein synthesis by regulating the efficiency whereby mRNAs are translated into proteins. Aberrant activation of mTORC1, as seen with TSC1/TSC2 loss, leads to hyperphosphorylation of key substrates controlling translation including eukaryotic translation initiation factor (eIF) 4E-binding proteins (4E-BPs [MIM: 602223 and MIM: 602224]) and p70 ribosomal protein S6 kinase 1 (S6K1 [MIM: 608938]). In the unphosphorylated state, i.e. when mTORC1 activity is low, 4E-BPs bind and sequester eIF4E [MIM: 133440]. This leads to reduced formation of the heterotrimeric eIF4F-complex, a key factor for initiation of translation, comprised of eIF4E, eIF4G [MIM: 600495] and eIF4A [MIM: 602641]. Activation of mTORC1 leads to phosphorylation of 4E-BPs and thereby release of eIF4E, which promotes eIF4F-complex formation and stimulates 5’ cap-dependent translation (Figure 1A). Phosphorylation of S6K protein affects protein synthesis and ribosome biogenesis through multiple mechanisms, including release of sequestered eIF4A via phosphorylation/inactivation of programmed cell death 4 (PDCD4 [MIM: 608610]) (Figure 1A) reviewed in ^11; 12^. Importantly, mTORC1 modulates translation, and thereby molds the proteome, in a transcript-selective fashion depending on features in the 5’ untranslated regions (UTRs) of target mRNAs. Three main classes of transcripts showing translation paralleling mTORC1 activity have been described: mRNAs with very short 5’UTRs, highly structured 5’UTRs and those with the 5′ terminal oligopyrimidine (5′ TOP) motif (reviewed in ^13^). Activation of mTORC1 is reported to be associated with suppressed translation of a set of mRNAs commonly harboring upstream open reading frames in their 5’UTRs ^14^. In addition, phosphorylation of eIF4E at a single conserved serine (S209) by mitogen-activated protein kinase-interacting kinase MNK1/2 [MIM: 606724 and MIM: 605069] also modulates translation in a transcript-selective fashion (Figure 1A) ^15; 16^.

**Figure 1.**
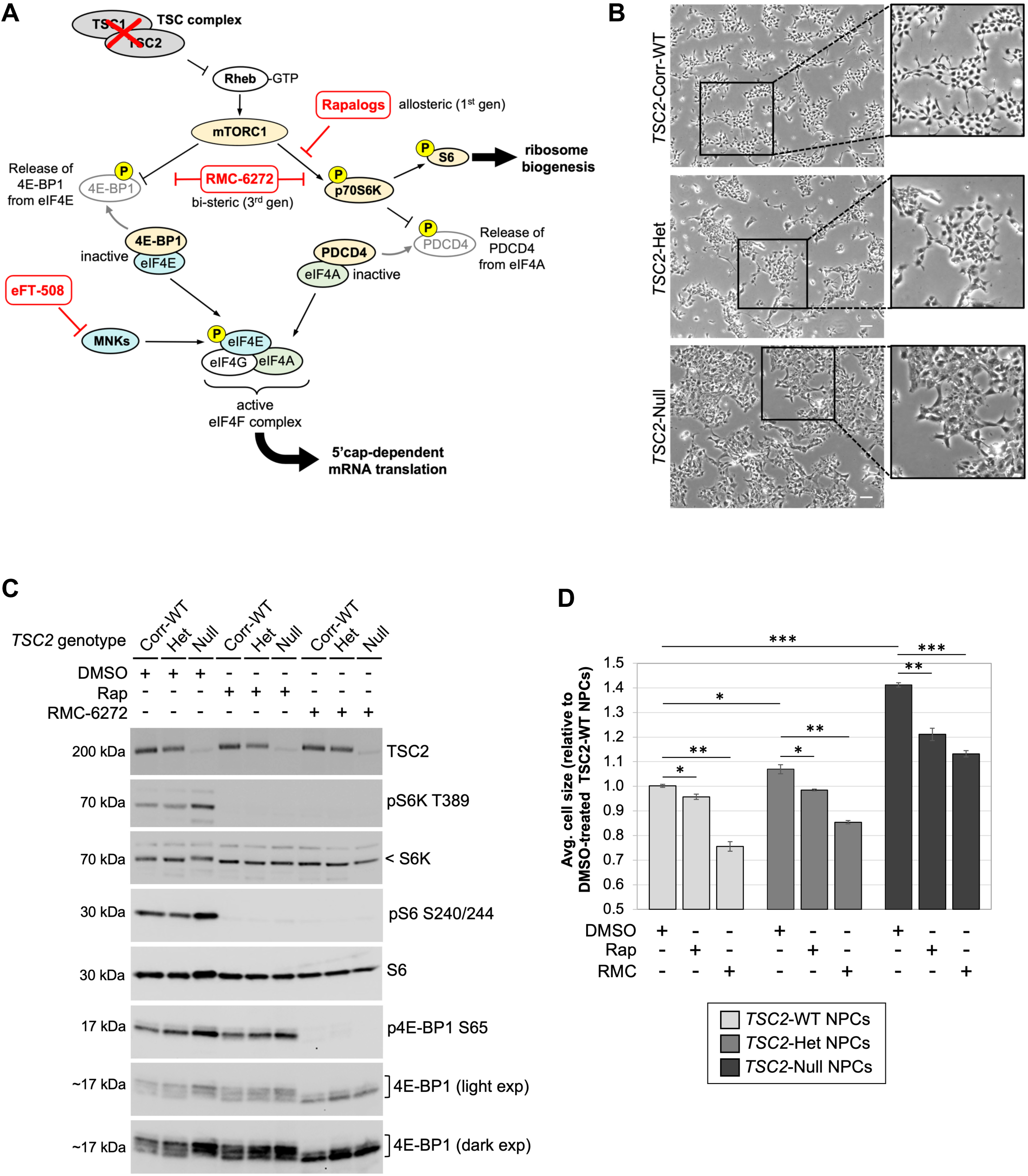
Inhibition of mTORC1 using RMC-6272 or rapamycin rescues increased cell size in *TSC2*-mutant NPCs. (A) Schematic representation of how mTORC1 signaling modulates mRNA translation. Inhibitors used to target the pathway axes are indicated in red. (B) Representative bright field images, including enlarged inset images (black boxed regions), are shown for *TSC2*-Het and -Null NPCs compared with *TSC2*-WT cells. Scale bar=100µm. (C) Immunoblotting of an isogenic set of *TSC2*-WT, -Het and -Null (cl. 39) NPCs, following 24 h treatment with DMSO (control), rapamycin (50nM) or RMC-6272 (10nM) are shown for TSC2/tuberin as well as mTORC1 pathway readouts pS6K T389, pS6 S240/244 and p4E-BP1 S65 along with respective total proteins. (D) Average cell size for *TSC2*-Het and -Null NPCs relative to *TSC2*-WT cells following a 48 h treatment with DMSO (control), rapamycin or RMC-6272 (same concentrations as in C) was quantitated, and data represents the mean (±SEM) from 3 independent experiments, each with triplicate technical replicates. P-values of comparisons (*p<0.05, **p<0.01, ***p<0.001) from Student’s t-test are indicated.

Although commonly used rapalogs efficiently suppress phosphorylation of S6K1, they only marginally affect 4E-BP1 phosphorylation in many cell types, which is consistent with rapalogs only partially inhibiting mTORC1 activity ^17–19^. This prompted the development of third-generation bi-steric mTORC1-selective inhibitors, Rapalinks, that link the mTORC1 affinity of rapamycin with a potent second-generation active-site mTOR kinase inhibitor, resulting in reduced phosphorylation of both S6K1 and 4E-BP1 ^20^. A more recent third-generation bi-steric mTORC1-selective inhibitor, RMC-6272 (a preclinical tool compound for its clinical counterpart RMC-5552), shows dramatically improved mTORC1 selectivity as compared to the Rapalink prototype while potently reducing phosphorylation of both S6K1 and 4E-BP1 (Figure 1A) ^21; 22^. Moreover, we recently demonstrated that RMC-6272 is superior to rapamycin in durably inhibiting the mTORC1-4E-BP1 axis and suppressing tumor growth in patient-derived *NF2*-deficient models ^23^. RMC-6272 has also shown strong anti-tumor activity as a single agent or when combined with other treatments in several preclinical cancer models ^21^, and the first-in-human study of RMC-5552 has offered preliminary data of anti-tumor activity at tolerated doses (NCT04774952 and Schram et al., 2023 https://doi.org/10.1158/1535-7163.TARG-23-C020). Additionally, recently developed eFT-508/tomivosertib is a potent and selective MNK1/2 inhibitor that effectively blocks phosphorylation of eIF4E (Figure 1A) ^24^ and is in phase 2 clinical trials for several cancer types (NCT02937675, NCT02605083 and NCT03258398).

While numerous global studies of mRNA levels (“transcriptomes”) in patient-derived neuronal models of ASD, TSC and other NDDs have been reported, studies of mRNA translation are largely lacking. Furthermore, the effects on mRNA translation from highly specific mTORC1 and MNK inhibitors such as RMC-6272 and eFT-508 in NDDs have never been explored. Our recent studies employing polysome-profiling to examine mRNA translation at a transcriptome-wide level in TSC1 patient-derived neural progenitor cells (NPCs) uncovered *TSC1*-associated transcript-selective alterations in translation that were largely insensitive to rapamycin. Interestingly, in confirmational studies using NanoString analysis, we observed that a small subset of selected mRNAs encoding proteins related to synaptic regulation or ASD were rapamycin-insensitive but showed sensitivity to RMC-6272. Moreover, unlike rapamycin, RMC-6272 also reversed some of the neurodevelopmental phenotypic defects in *TSC1*-Null NPCs ^25^. These findings suggest a key role of mTORC1-4E-BP1-eIF4E axis-dependent modulation of mRNA translation in TSC. Here, we have further investigated the role of this axis in regulating mRNA translation using an isogenic set of CRISPR-modified TSC2 patient-derived NPCs. Our results reveal a strong overlap in dysregulated translation observed in *TSC1*-Null and *TSC2*-Null NPCs. Surprisingly, genes previously associated with ASD and NDD were translationally suppressed in *TSC2*-Null NPCs, and their translation was reversed upon RMC-6272 or eFT-508 treatment. These results establish the importance of mTORC1-eIF4F- and MNK-eIF4E-mediated mRNA translation in TSC, ASD and NDDs and lay the groundwork for evaluating drugs in clinical development that target these pathways as a treatment strategy for TAND as well as ASD/NDD.

## MATERIAL AND METHODS

### Human induced pluripotent stem cell derivation

We previously reported collection of post-mortem subungal fibroma (SUF) tissue from a female sporadic TSC2 patient, which was obtained through the pathology department and approved by the Institutional Review Board at Massachusetts General Hospital. Detailed genetic analysis of the patient SUF tissue, including exon-by-exon scanning of the entire coding sequence of *TSC2* isoform 1 (GenBank: NM_000548), identified a single heterozygous (*TSC2*-Het) truncating nonsense mutation in *TSC2* exon 34 (originally reported as coding exon 33; 4207delG, D1400fs>1410X). Further analysis of the SUF tissue for tumor clonality and loss-of-heterozygosity (LOH), using multiple polymorphic chromosome 16 or *TSC2* intragenic markers, revealed a clonal tumor with no second-hit mutations or LOH ^26^. For culturing of primary *TSC2*-Het fibroblasts, at the time of collection SUF tissue was rinsed twice in Hank’s Balanced Salt Solution (Sigma) followed by mincing in high glucose DMEM (Gibco) containing 15% fetal bovine serum (FBS; Sigma), 0.5 µg/ml fungizone (Life Technologies), 50 g/ml gentamicin (Life Technologies), and 1x penicillin-streptomycin/L-glutamine (Cellgro). The tissue was then digested in the same medium as above (without gentamycin) containing ∼5 units collagenase III (Fisher Scientific), for 30 min at 37^0^C. Digested tissue was then pelleted by centrifugation, resuspended in high glucose DMEM (containing 15%FBS, 1x penicillin-streptomycin/L-glutamine), and the resulting cultured fibroblast line was then expanded and cryopreserved in liquid nitrogen. Following revival of cryopreserved cells, *TSC2*-Het fibroblasts were reprogrammed using synthetic modified mRNA-based methods to obtain induced pluripotent stem cells (iPSCs) as described ^27; 28^. Briefly, cells were transfected by nucleo-electroporation (Amaxa Nucleofector I) with *in vitro* transcribed mRNAs encoding *Oct4, Sox2, Klf4, c-Myc* and *Lin28* (Stemgent). After single clone picking, iPSC colonies were cultured in feeder-free conditions in Essential 8 medium (ThermoFisher) on plates coated with Geltrex (Gibco) using the “thin gel method” according to the manufacturer’s instructions. Colonies were passaged every 4-6 days with daily medium change. Sanger sequencing confirmed the original heterozygous mutation in *TSC2*-Het iPSCs (see details below for *TSC2* exon 34 PCR and sequencing).

### CRISPR/Cas9-mediated generation of isogenic iPSC lines

To generate both the corrected-wildtype (WT) and Null isogenic iPSC lines, CRISPR/Cas9 genome editing was performed employing parental *TSC2*-Het iPSCs. To achieve gene knock-in to correct the germline *TSC2* mutation, we followed the same cloning and transfection strategy as we previously reported for *TSC1* corrected-WT iPSCs ^29^. Briefly, using web-based CRISPR-related tools developed by GenScript (https://www.genscript.com), we designed a single guide RNA (sgRNA): 5’-GGACATCCTCGGGGACCCTG-3’ targeting *TSC2* exon 34 (coding exon 33) along with an asymmetric donor sequence: 5’-CCGAGGCCGACCAGGCAGCACTTTCCCCGTCCAGGGTCCCTGACTGTGACCGGGCCTTAACCTCAGGGCTCAGCCGGCCCACGTCGGCCTTGTCCCCAGGGTCCCCGAGGATGTCCTGCAGAGTCTGCAG-3’ for homology-directed repair. Colonies were manually isolated, and genomic DNA was extracted from a portion of each colony followed by PCR amplification using primers TSC2-ex33F: 5’-CTGACAGGGGTTCTCTTTGG-3’ and TSC2-ex33R: 5’-TCCAGGGTCCCTGACTGTGA-3’. Screening by Sanger sequencing, using a nested primer TSC2-ex33-seqF: 5’-TCGTCCTCAGTCTCCAGCC-3’, identified one clone with successful correction back to WT. For generating the *TSC2*-Null iPSC lines, we used a service provided by an independent company (Alstem Cell Advancements, Richmond, CA). Briefly, *TSC2*-Het iPSCs were co-transfected by nucleo-electroporation (Amaxa Nucleofector I) with a pair of sgRNAs targeting *TSC2* exon 2 (coding exon 1) including sgRNA-1: 5’-TGTTGGGACTGGGAACACCG-3’ and sgRNA-2: 5’-ACTCACTCTCAGTATTTCCG-3’. A total of 50 single clones were screened by PCR amplification and Sanger sequencing using primers TSC2-F2: 5’-CCATGGCCAAACCAACAAG-3’ and TSC2-R2: 5’-ACCTGGTGCAAGACCAAA-3’, with 3 homozygous *TSC2*-Null clones identified, all of which harbored the same 59 bp deletion in *TSC2* exon 2 resulting in a frameshift and premature protein truncation.

### Differentiation of iPSCs into NPCs

To generate NPCs, *TSC2*-iPSCs including the parental *TSC2*-Het line, the single *TSC2*-WT clone and two of the three *TSC2*-Null clones (clones 5 and 39) were differentiated using the directed monolayer differentiation protocol as we previously reported for TSC1-NPCs ^29;30^. Briefly, iPSCs expressing the pluripotency marker TRA-1-60 were sorted and enriched using magnetic-activated cell separation (MACS) technology (Miltenyi Biotec). Cells were cultured in neural induction medium (neurobasal medium supplemented with 1x neural induction supplement (ThermoFisher)) for 7-9 days, followed by a second sorting for cells expressing polysialylated-neural cell adhesion molecule (PSA-NCAM). The PSC-NCAM-positive (+) cells then underwent a double-sorting to enrich for NPCs representing CD271(-) /CD133(+) cells. NPCs were then cultured in neural expansion medium (1:1 neurobasal medium:advanced DMEM/F12, 1x neural induction supplements (ThermoFisher) and 1x penicillin/streptomycin (Corning)). MACS sorting was performed using microbead kits for TRA-1-60, PSA-NCAM, CD271 and CD133 (Miltenyi Biotec).

### Immunocytochemistry

To confirm successful derivation of NPCs, immunofluorescent staining was carried out using NPC markers Nestin (EMD Millipore) and Sox2 (R&D Systems) antibodies as previously described ^29; 30^. Briefly, cells were fixed with 4% paraformaldehyde (PFA; Electron Microscopy Sciences) and then permeabilized and blocked in a single step using PBS containing 4% normal goat serum (NGS)/0.1% Triton-X-100. Primary antibodies were diluted into PBS containing 2% NGS/0.1% Triton-X-100. DAPI (Invitrogen) was used to stain nuclei at 5 μg/ml. Coverslips were mounted in ProLong Gold antifade Mountant (Invitrogen) and images were captured using a Nikon Eclipse TE2000-U microscope and the NIS-Element BR 3.2 imaging software.

### Immunoblot analysis

Cells were lysed in RIPA lysis buffer (50 mM Tris (pH7.5), 150 mM NaCl, 0.1% SDS, 0.5% sodium deoxycholate, 1% NP-40, 1x HALT phosphatase inhibitor cocktail (Pierce) and 1x protease inhibitor cocktail (Sigma)) as previously described ^31; 32^. Protein lysates were resolved on 4-12% or 10-20% Novex WedgeWell Tris-glycine gels (Invitrogen), transferred to 0.45 μm pore nitrocellulose (BioRad) and then incubated with primary antibodies to detect TSC2, pS6K T389, S6K, pS6 S240/244, S6, p4E-BP1 S65, 4E-BP1, peIF4E S209, and eIF4E (Cell Signaling Technologies); β-actin (Santa Cruz Biotechnology) or GAPDH (EMD Millipore). All immunoblotting data shown are representative of at least 3 biological replicates.

### Cell Size and proliferation

Cell size and proliferation were determined for *TSC2*-WT, Het and Null NPCs treated for 72 h with 50 nM rapamycin (MilliporeSigma), 10 nM RMC-6272 (generously provided by Revolution Medicines Inc.) or DMSO as vehicle control. For both cell size and proliferation, NPCs were analyzed by trypan blue exclusion using the Countess II automated cell counter (ThermoFisher) as previously described ^25; 29^. Briefly, cells were detached using Accutase enzyme detachment medium (ThermoFisher), and for each sample, 10 μl of suspended cells were combined with 10 μl of trypan blue, and then 10 μl of the resulting mixture was assessed using the Countess II automated cell counter (ThermoFisher), according to the manufacturer’s instructions. Quantification of viable and dead cells was performed along with size of viable cells. For proliferation, counting of viable cells collected at days (D) 1, 2, 3 and 4 after seeding was performed for cells treated with 50 nM rapamycin, 10 nM RMC-6272 or DMSO on D1. All data collected using the Countess II cell counter were analyzed from 3 independent experiments (3 technical replicates each). Proliferation was further assessed using flow cytometry employing the proliferation marker Ki-67 as previously described ^25^. Briefly, *TSC2*-WT, Het and Null NPCs were seeded onto Geltrex-coated plates followed by treatment with 50 nM rapamycin, 10 nM RMC-6272, 50 nM eFT-508 (Selleckchem) or DMSO as vehicle control for 72 h (two biological replicates). To assess Ki-67 proliferation status, NPCs were fixed with 4% PFA, permeabilized with ice cold 100% methanol to a final concentration of 90% with gentle vortexing and stained using AlexaFluor (AF) 488-conjugated Ki-67 monoclonal antibody or AF488-conjugated IgG as an isotype control (Cell Signaling Technologies). Data were acquired from ∼1×10^4^ cells using a BD LSR II Flow Cytometer (BD Biosciences), and analysis was carried out using FlowJo 10.8.1 (FlowJo LLC). Percentages of Ki-67 positive cells (acquired through FITC channel) were determined by gating with respect to AF488-IgG control.

### Neurite outgrowth assay

NPCs were assessed for neurite outgrowth as previously described ^29^. Briefly, NPCs were seeded on coverslips coated with poly-D-lysine (0.1 mg/ml, Sigma) and fibronectin (5 μg/ml, Corning) in growth factor-depleted neural expansion medium containing 1:1 neurobasal medium:advanced DMEM/F12 (ThermoFisher), 0.3x neural induction supplements (ThermoFisher) and 1x penicillin/streptomycin (Corning). Cells were treated with 50 nM rapamycin, 10 nM RMC-6272 or DMSO vehicle control for 48 h followed by fixation with 4% PFA prior to immunostaining. Three independent experiments were performed where images were acquired from 8 non-overlapping fields and analyzed using HCA-Vision software V.2.2.0 (CSIRO, Canberra, Australia) ^33^. Images were acquired using a Nikon Eclipse TE2000-U microscope and NIS-Element BR 3.2 imaging software.

### Polysome fractionation and RNA sequencing

Lysate were prepared for polysome profiling from three biological replicates as previously described with minor modifications ^34^. Briefly, TSC2-NPC lines were seeded at ∼50,000 cells/cm^2^, with a total of ∼2.4×10^7^ cells seeded per replicate for three drug treatment conditions, including 10 nM RMC-6272, 50 nM eFT-508 or DMSO control. The day after seeding, cells were treated for 2 h, rinsed with 1x PBS containing 100 μg/ml cycloheximide (CHX) (Sigma), harvested by scraping on ice in PBS/CHX and pelleted by centrifugation at 300g for 10 min at + 4C. Cell pellets were first resuspended in hypotonic lysis buffer containing 5 mM Tris-HCl (pH 7.5), 2.5 mM MgCl_2_ and 1.5 mM KCl, followed by addition of DTT (2 mM final), CHX (100 µg/ml final), sodium deoxycholate (0.5% final) and Triton X-100 (0.5% final). Lysates were then vortexed for 4 sec, cleared by spinning for 2 min at 13,000 rpm and quickly frozen on dry ice. When ready for processing, lysates were quickly thawed at 37°C and loaded onto a 10 - 50% sucrose gradient, followed by centrifugation for 2 h and 15 min at 35,000 rpm in a SW41 rotor using a Sorvall Discovery 90SE centrifuge. The gradients were fractionated on a Teledyne ISCO Foxy R1 apparatus while monitoring the OD254nm. Fractions corresponding to mRNA associated with more than two ribosomes were pooled and the RNA extracted using TRIzol (ThermoFisher) according to the manufacturer’s protocol. 10% of the lysate was removed from the extract prior to loading on the sucrose gradient, extracted using TRIzol, and denoted as total RNA. RNA sequencing libraries were prepared from the resulting samples (both polysome-associated and total RNA) using Illumina v2.5 Kits and sequenced on an Illumina NextSeq 500 at the Canada’s Michael Smith Genome Sciences Centre (BC Cancer Research Institute, Vancouver, Canada).

### RNA sequencing data preprocessing and quality control

Adapters were removed from RNA sequencing reads using bbduk.sh from the BBTools software suite [https://www.osti.gov/servlets/purl/1241166] (v. 36.59 with settings: k = 13, ktrim = n, useshortkmers = t, mink = 5, qtrim = t, trimq = 10, minlength = 20). Paired-end reads were aligned to hg38 using HISAT2 ^35^ (ver 2.1.0 with settings: --no-mixed, --no-discordant), and uniquely mapped reads were quantified using the featureCounts function from the RSubreads package (v.2.6.4) ^36^ with RefSeq gene definitions and default settings. Polysome-profiling data from the previous study of *TSC1*-dependent translation^25^ were quantified using the same gene definitions to allow a direct comparison to the herein generated dataset. Dataset reproducibility was assessed using Principal Component Analysis using genes within the highest quartile of standard deviations quantified across all samples as input.

### Analysis of gene expression alterations using anota2seq

Gene expression quantified from polysome-associated and total mRNA was analyzed using the anota2seq algorithm (v. 1.20.0) ^37^. Batch effects were accounted for in the models by including sample replicates using the “batchVec” argument in anota2seq. Default threshold settings were used [settings minSlopeTranslation = -1, maxSlopeTranslation = 2, minSlopeBuffering = -2, maxSlopeBuffering = 1, deltaPT = deltaP = deltaTP = deltaT = log2(1.2), minEff=log2(1.5)] with a significance threshold maxRvmPAdj = 0.15 (corresponding to a false discovery rate [FDR] < 0.15). Transcripts were classified into three modes of regulation assessed by anota2seq (changes in mRNA abundance, changes in translation, and translational offsetting) using the “anota2seqRegModes’ function.

### Analysis of gene signatures using empirical cumulative distribution functions

Empirical cumulative distribution functions (ECDFs) of log2 fold changes for polysome-associated and total mRNA were plotted for gene signatures including ASD and NDD genes ^38^, eIF4E overexpression ^39^, mTOR-sensitive genes ^40^ obtained from previous publications or, to cross-compare datasets, genes that were found to be translationally regulated in either dataset. The Wilcoxon rank-sum test was used to determine whether there was a significant shift between each signature and the background (i.e. genes not within the signature).

### Gene set enrichment analysis

Gene Ontology (GO) enrichment analysis was performed using genes classified as regulated via translation by anota2seq (differentially translated genes, “DTGs”) and annotation from GO biological process (GO:BP) terms retrieved from MSigDB database (v. 2023.2.Hs) ^41^ using one-tailed Fisher’s exact test. In this analysis, only GO:BP terms with at least 10 genes within the background set (all detected genes analyzed by anoat2seq) of a particular comparison were considered. DTG datasets for GO analysis included *TSC2*-Null vs WT NPCs, *TSC2*-Null NPCs treated with RMC-6272 vs DMSO control, *TSC2*-Null NPCs treated with eFT-508 vs DMSO control, pairwise comparison between *TSC2*-Null NPCs treated with RMC-6272 or eFT-508 as well as pairwise comparisons between *TSC2*-Null vs WT NPCs and our previously published dataset comparing *TSC1*-Null vs WT NPCs ^25^. In addition, genes were ranked based on fold-changes for translation determined by anota2seq analysis for a given comparison. The analysis utilized ASD-, NDD- and EPI-associated gene sets. This was conducted using the fgsea package (v.1.24.0, https://doi.org/10.18129/B9.bioc.fgsea) within the R environment. For each gene set, the Normalized enrichment score (NES) and FDR were calculated.

Enrichment analysis for selected DTGs identified by anota2seq and target gene sets, including ASD-, NDD-, EPI-associated genes ^38; 41;42^ and eIF4E-sensitive genes ^39^, was performed using one-tailed Fisher’s exact test. Statistical significance of overlaps between the two DTG datasets was also assessed using one-tailed Fisher’s exact test. Here, all anota2seq analyzed genes in a single comparison, or intersection of anota2seq analyzed genes in paired comparisons, constituted the background sets. These enrichment analyses were performed in the R statistical software environment (version 3.4.3) (https://www.R-project.org/) using custom scripts.

### Validation of differential translation using RT-qPCR

For validation, polysome fractionations were collected from *TSC2*-Null or *TSC2*-WT NPCs (three biological replicates), and RNA was isolated as described for NPCs above. cDNA was prepared using M-MuLV Reverse Transcriptase (New England Biolabs) and oligo(dT)20 primers using the manufacturer’s recommendations. RT-qPCRs were performed with SsoFast EvaGreen Supermix using the CFX96 PCR system (Bio-Rad). Primers for RT-qPCR are detailed in Table S1. The level of each mRNA was normalized to β-actin (*ACTB* [MIM: 102630]) and Phosphoglycerate Kinase 1 (*PGK1* [MIM: 311800]) using the comparative CT method and compared across conditions as indicated in the figure legend.

### Statistical analyses

Methods for statistics are described above, or in corresponding figure legends.

## RESULTS

### Establishment of an isogenic set of TSC2 patient-derived neural progenitor *cells*

We generated an isogenic set of TSC2 patient-derived NPCs using a similar approach as we described previously for TSC1 patient-derived NPCs ^25; 29^. Primary fibroblasts cultured from post-mortem SUF tissue were found to harbor a germline heterozygous 1 bp deletion in *TSC2* exon 34 (originally reported as coding exon 33) with no loss-of-heterozygosity ^26^. Induced pluripotent stem cells (iPSCs) were generated from the fibroblasts, followed by CRISPR-based genome editing that demonstrated (i) correction of the heterozygous mutation back to wildtype, or (ii) knock out of the second allele due to a 59 bp deletion in *TSC2* exon 2 (coding exon 1). This resulted in an isogenic set of iPSCs representing the parental *TSC2*-Het line, a single clone of *TSC2*-corrected (*TSC2*-WT) and three clones of *TSC2*-Null iPSCs. We then selected *TSC2*-WT, Het and two Null iPSC clones (clones 5 and 39) for microbead sorting of neural cell surface protein markers to obtain NPCs as previously described ^29^. Immunostaining for neural markers Nestin and Sox2 confirmed successful NPC derivation (Figure S1A). Loss of TSC2 protein expression was confirmed in *TSC2*-Null clones when compared with the *TSC2*-WT (Figure S1B). We also observed increased phosphorylation of mTORC1 targets p70S6K (pS6K T389), pS6 S240/244 and p4E-BP1 S65 in both *TSC2*-Null NPC clones compared with *TSC2*-WT NPCs, consistent with activation of mTORC1 signaling, while *TSC2*-Het NPCs revealed only minimal activation of mTORC1, which is in agreement with several reports that employed human iPSC models of TSC ^43–46^ (Figure S1B). *TSC2*-Null NPCs demonstrated an enlarged cell phenotype compared with *TSC2*-WT cells (Figure 1B, D), which is a hallmark of mTORC1 activation. *TSC2*-Het NPCS exhibited an intermediate cell size phenotype (Figure 1B, D). Treatment with rapamycin or RMC-6272 attenuated phosphorylation of S6K at T389, along with its downstream substrate ribosomal protein S6 (pS6 S240/244) and rescued the enlarged cell size (Figure 1C, D). As expected, rapamycin showed only minimal effects on phosphorylation of 4E-BP1 while RMC-6272 demonstrated complete inhibition (Figure 1C). These results are consistent with our observations in TSC1-NPCs ^25^ and with previous studies showing that the mTORC1-S6K signaling axis controls cell size ^47^.

### Rescue of early neurodevelopmental phenotypes by RMC-6272, but not rapamycin

We previously demonstrated that defects in early neurodevelopmental phenotypes in *TSC1*-Null NPCs, including increases in proliferation rate and neurite outgrowth, could be rescued by RMC-6272 treatment, but not rapamycin. This suggests dependence, at least in part, on the mTORC1-4E-BP1 signaling axis ^25; 29^. Here, we assessed proliferation by viable cell count in TSC2*-*NPCs over the course of four days (D1-4) after seeding, also comprising a 72 h treatment with 50 nM rapamycin, 10 nM RMC-6272 or DMSO control initiated at D1. We observed genotype-dependent changes in cell numbers in vehicle-treated cells whereby *TSC2*-Null NPCs showed significantly increased proliferation relative to *TSC2*-WT NPCs with *TSC2*-Het also showing augmented proliferation as compared to *TSC2*-WT NPCs (Figure 2A, left graph region). When compared with DMSO treatment, rapamycin showed no significant effect in *TSC2*-WT, -Het or -Null NPCs (Figure 2A, center compared with left graph regions); whereas RMC-6272 demonstrated significant reduction by D3 in all three NPC genotypes, with the maximal decrease observed in *TSC2*-Null NPCs (Figure 2A, right compared with left graph regions). Consistent with this, flow cytometry-based analysis showed a reduction in the percentage of S phase-positive cells (as indicated by Ki-67 positive staining) in RMC-6272 vs DMSO treated *TSC2*-WT (50.9% vs 62.7%), *TSC2*-Het (56.2% vs 64.9%) and *TSC2*-Null (53.7% vs 68.9%) NPCs. In contrast, similar percentages of Ki-67 positivity were found for rapamycin or DMSO treatment (Figure S2A). Moreover, treatment with eFT-508/tomivosertib, a MNK1/2 inhibitor, which attenuates phosphorylation of eIF4E at S209 (peIF4E S209) but not 4E-BP1 (p4E-BP1 S65) did not alter Ki-67 positivity in *TSC2*-Null NPCs (Figure S2A, B).

**Figure 2.**
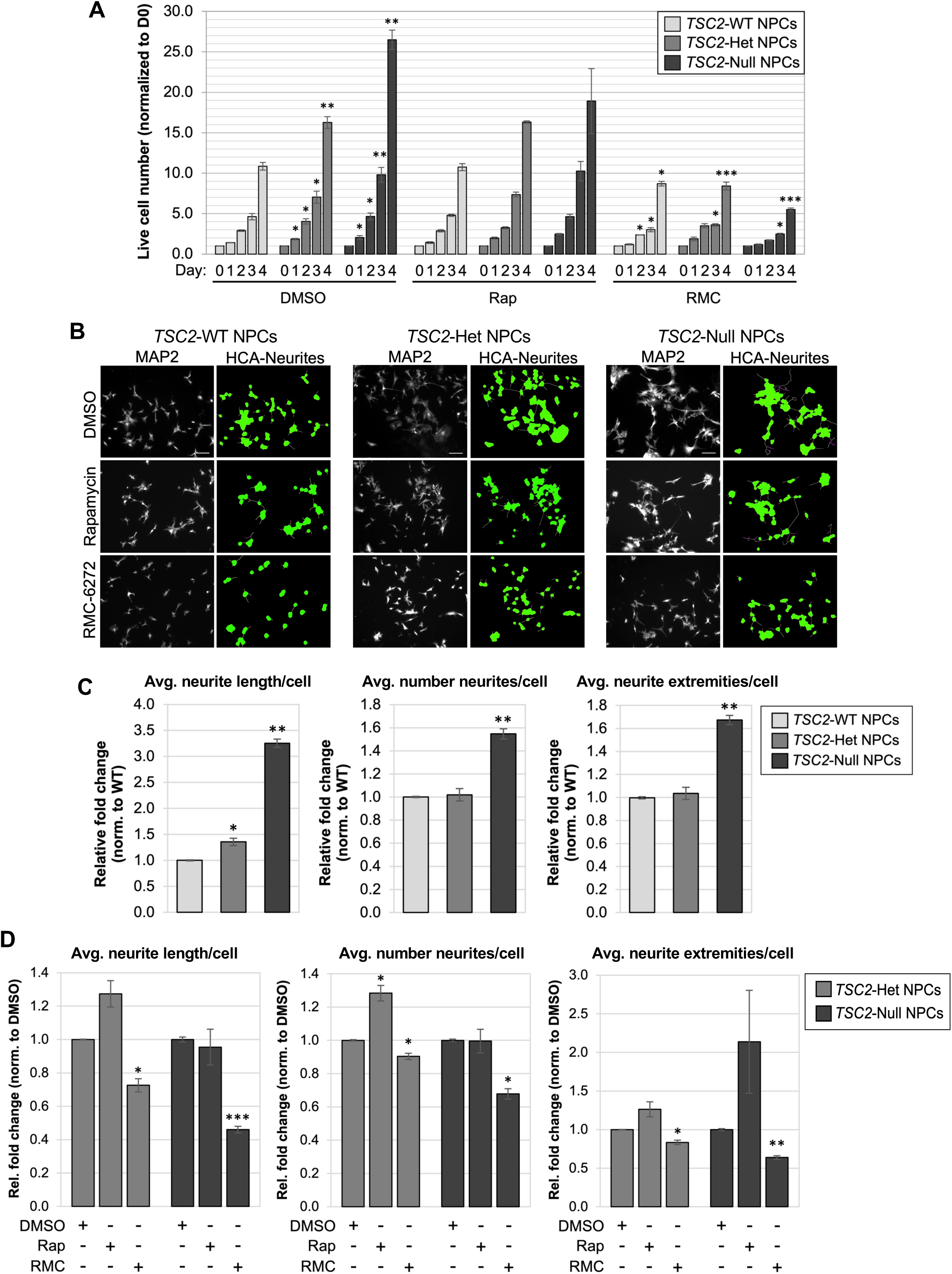
Altered neurodevelopmental phenotypes in *TSC2* mutant NPCs are rescued by RMC-6272, but not rapamycin. (A) Proliferation by live cell counting (trypan blue exclusion) was carried out over days D1-4 in *TSC2*-WT, -Het and -Null NPCs under control treatment (left graph region), or treatment initiated at D1 with rapamycin (50 nM, middle graph region) or RMC-6272 (10 nM, right graph region). Quantitation for each timepoint is normalized to cell seeding number (D0=1). Data represent the mean (±SEM) from 3 independent experiments (3 replicates/cell line/experiment). Student’s t-test p-values (*p<0.05, **p<0.01, ***p<0.001) are indicated in DMSO-treated NPCs (left graph region) for increasing cell numbers in *TSC2*-Het and Null NPCs at each timepoint (D1-4) relative to *TSC2-*WT NPCs at the respective timepoint. For drug-treated NPCs (middle and right graph regions), within each genotype, p-values are indicated for changes in cell numbers relative to DMSO controls at the corresponding timepoint. (B) Representative images of immunofluorescence staining in NPCs using neuronal marker MAP2 (gray), performed 48 h after initiating nerite outgrowth, are shown along with traced images generated employing HCA-Vision image quantitation software (green), which traces the cell body along branching layers of the neurite arborization. Scale bar = 100 μm. (C and D) Bar plots indicate average neurite length (left panel), average neurite number (center) and extremities (right) per cell for *TSC2*-Het and -Null NPCs relative to *TSC2*-WT NPCs under control conditions (C) or for rapamycin (50 nM) or RMC-6272 (10 nM) treatment in *TSC2-*Het and -Null cells relative to DMSO control (D) . Data represents mean (±SEM) from 3 independent experiments (8 non-overlapping image fields/cell line/experiment). P-values from Student’s t-test (*p<0.05, **p<0.01, ***p<0.001) are shown.

We next assessed neurite outgrowth in isogenic TSC2-NPCs using MAP2 immunostaining and HCA-Vision image analysis software to quantitate three parameters (as an average per cell): (i) length of neurite outgrowth, (ii) number of neurite roots and (iii) number of extremities (terminal endpoint of primary and branched outgrowths). In TSC2-NPCs treated for 48 h with DMSO, a significant increase in neurite length/cell of ∼3.2-fold in *TSC2*-Null and an intermediate increase of ∼1.4-fold in *TSC2*-Het was observed relative to *TSC2*-WT NPCs (Figure 2B, C). *TSC2*-Null NPCs also revealed a significant increase in neurite number/cell (∼1.5-fold) and extremities/cell (∼1.7-fold) relative to *TSC2*-WT NPCs (Figure 2B, C). Upon RMC-6272 treatment, we observed decreases in neurite length, number and extremities of 0.73, 0.90 and 0.83, respectively, for *TSC2*-Het NPCs, and 0.46, 0.68 and 0.64, respectively, for *TSC2*-Null NPCs (Figure 2D). Similar to the proliferation assays, rapamycin failed to rescue neurite-related phenotypes, and in fact showed an opposite trend in several parameters (Figure 2D). Taken together, these data support that the third-generation mTORC1 inhibitor RMC-6272 more effectively reverses the early neurodevelopmental phenotypes observed in TSC2-NPCs, and that this effect is dependent on a more potent inhibition of mTORC1-4E-BP signaling.

### Abundant mRNA translation alterations in *TSC2*-Null NPCs largely overlap with those observed in *TSC1*-Null NPCs

Our findings above demonstrate activation of mTORC1 signaling and increases in proliferation and neurite outgrowth in *TSC2*-Null NPCs, compared with WT cells. As mRNA translation is a major downstream target of mTORC1, it is necessary to study how both mRNA levels and translation are affected by *TSC2* loss to disentangle gene expression-dependent effects on cellular phenotypes. We therefore applied polysome-profiling in *TSC2*-Null and *TSC2*-WT NPCs. Cytoplasmic mRNA was fractionated based on ribosome-association and the resulting polysome-associated (herein mRNA associated with >2 ribosomes) and total cytoplasmic mRNA pools were then quantified using RNA sequencing followed by anota2seq analysis ^37; 48^. This allowed identification of *TSC2*-associated alterations in (i) “translation”, defined as polysome-associated mRNA alterations not paralleled by corresponding changes in total mRNA levels; (ii) “abundance”, representing congruent modulation of total and polysome-associated mRNA; and (iii) “offsetting”, defined as alterations in total mRNA not paralleled by corresponding changes in polysome-associated mRNA. This analysis revealed numerous *TSC2*-associated alterations in mRNA translation visualized using a scatterplot comparing *TSC2*-dependent changes in polysome-associated and total mRNA (Figure 3A; Table S4) and densities of p-values and false discovery rates (FDRs) obtained from anota2seq analysis (Figure 3B). These alterations in translation occurred alongside less abundant changes in mRNA levels and essentially in absence of translational offsetting (Figure 3A). Although the analysis is expected to be hampered by the large number of regulated genes, we used gene ontology analysis to identify cellular functions targeted by these changes in translation. This revealed that translationally suppressed transcripts encode proteins enriched for annotation to cellular functions including protein DNA complex organization, chromatin remodeling, RNA metabolism, and cell cycle while those involved in a variety of metabolic processes were translationally activated (Figure 3C).

**Figure 3.**
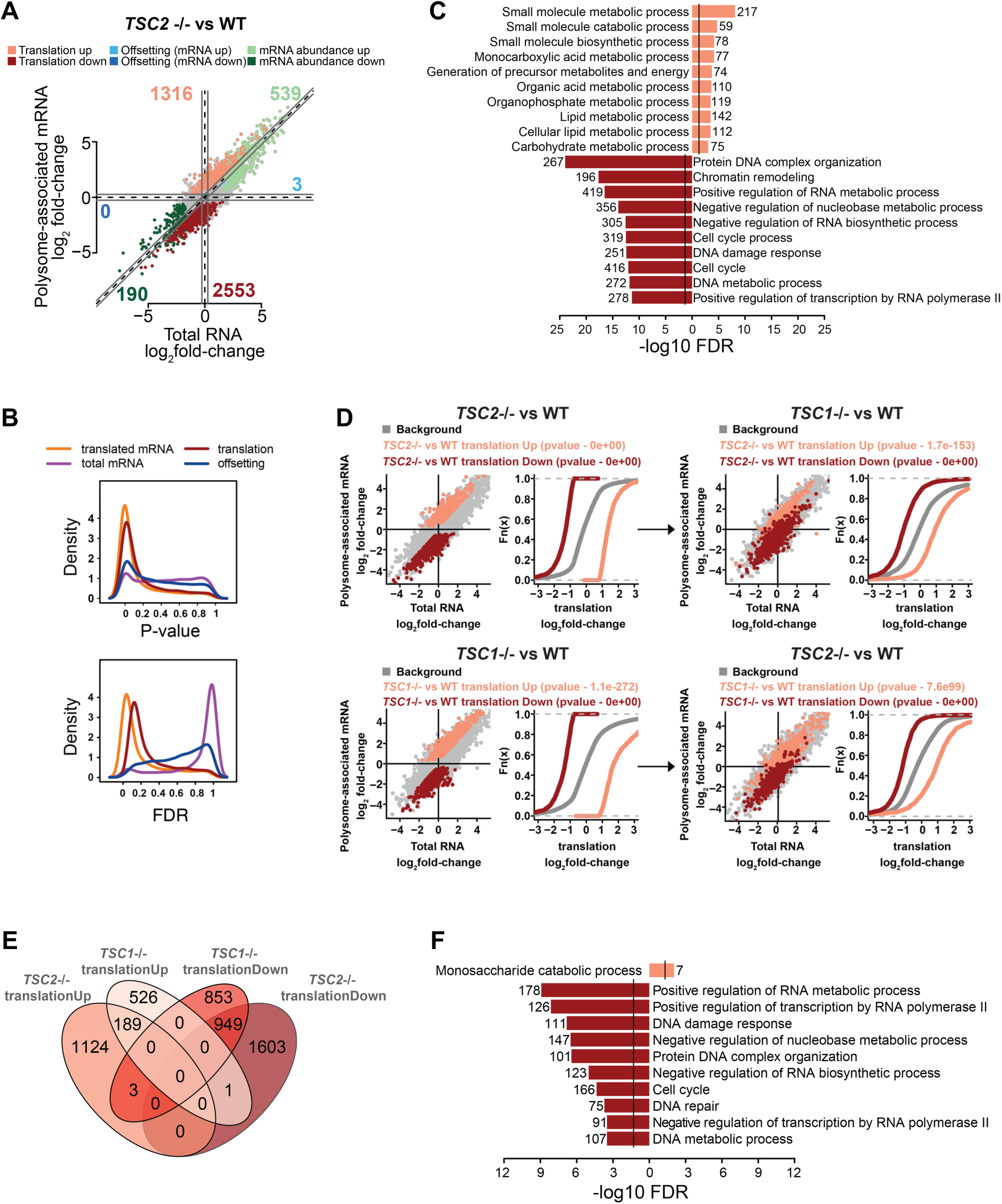
Extensive remodeling of mRNA translation in *TSC2*-Null NPCs aligns with those observed following loss of *TSC1*. (A) Scatterplot of polysome-associated mRNA vs total mRNA log2 fold changes between *TSC2*-Null (-/-) and WT NPCs. Genes are colored according to their mode of regulation as identified by the anota2seq analysis (FDR < 0.15). The number of mRNAs regulated via translation, mRNA abundance or offsetting is indicated. (B) Kernel densities of p-values and FDRs (from anota2seq analysis) for the comparison of *TSC2*-Null vs WT NPCs. Densities are shown for analysis of polysome-associated RNA (translated mRNA), total mRNA, translation and offsetting. A shift of the density towards low p-values/FDRs indicates a higher frequency of changes. (C) Enrichment of gene ontology annotation for biological processes among genes that undergo significant changes in translation (light red – activated translation, dark red – suppressed translation) in *TSC2*-Null vs WT NPCs. The ten most significant processes are shown. (D) Scatterplots from anota2seq analysis of *TSC2*-Null vs WT NPCs (top left and bottom right panels) and *TSC1*-Null vs WT NPCs (top right and bottom left panels). Transcripts identified as regulated by translation in anota2seq analysis in the *TSC2*-Null vs WT NPC (top left and right panels) or *TSC1*-Null vs WT NPC (bottom left and right panels) comparisons are indicated. Also shown are empirical cumulative distribution functions of the log2 fold changes in translation for the corresponding gene sets. Grey dots and lines correspond to the background (i.e. not in gene sets). Significant shifts of gene sets, relative to background, were determined using Wilcoxon rank-sum test and the resulting p-values are shown. (E) Venn diagram comparing translationally regulated genes identified in anota2seq analysis following loss of *TSC1* or *TSC2*. (F) Enrichments of gene ontology annotation for biological processes among genes identified as showing consistent regulation in both *TSC1*-Null and *TSC2*-Null NPCs relative to their controls (same color codes as in panel C).

To assess how these *TSC2*-associated alterations in mRNA translation compare to our previously described *TSC1*-associated changes, we looked for differences in the response to loss of *TSC1* vs *TSC2* relative to their respective control conditions (i.e. the interaction effect) using an anota2seq-like approaches we described previously ^49^. Strikingly, only one gene showed a statistically significant difference in translation response upon *TSC1* vs *TSC2* loss (at FDR < 0.15), and this analysis therefore argues for largely overlapping effects on mRNA translation. To further assess this, we generated gene signatures of transcripts that were translationally up or downregulated in *TSC1*-Null or *TSC2*-Null NPCs relative to their controls and visualized how these gene sets were regulated across the two models. This revealed that transcripts whose translation was activated or suppressed in *TSC2*-Null NPCs showed similar regulation in *TSC1*-Null NPCs and, vice versa, transcripts whose translation was activated or suppressed in *TSC1*-Null NPCs showed similar regulation in *TSC2*-Null NPCs (Figure 3D). Yet, there were differences in mRNA level changes for these subsets (compare Figure 3D left and right plots) that, together with expected threshold-effects, resulted in a smaller set of 949 translationally suppressed and 189 translationally activated transcripts that passed threshold applied in anota2seq for both comparisons (Figure 3E). As expected, these shared genes were enriched in similar gene ontology categories as those identified upon *TSC2* loss (Figure 3F). In aggregate, these results indicate extensive remodeling of mRNA translation in *TSC2*-Null NPCs largely recapitulating that observed in *TSC1*-Null NPCs.

### Suppressed translation of mRNAs encoding proteins associated with ASD and NDD upon *TSC2* loss

To explore the relationship between altered gene expression associated with *TSC2* loss and patient phenotypes we reasoned that *TSC2* loss may affect translation of mRNAs encoding proteins implicated in non-syndromic autism spectrum disorder (ASD), general neurodevelopmental disorders (NDD) and/or epilepsy. To assess this, we utilized data from recent studies that identified high confidence *de novo* mutations in large cohorts of individuals diagnosed with ASD, NDD as well as epilepsy. For ASD/NDD-related genes, we used datasets reported by Fu and colleagues, including 72 genes associated with ASD (ASD72) and 373 genes associated with NDD (NDD373) identified at a stringent FDR ≤ 0.001 threshold, as well as 185 ASD-associated genes (ASD185) and 664 NDD-associated genes (NDD664) at reduced stringency (FDR ≤ 0.05) ^38^. For epilepsy-related genes, we used genes categorized by the Epi25 Collaborative (http://epi-25.org/genes-in-epilepsy) and reported by Chen and colleagues ^42^ that consists of 83 genes identified at p ≤ 0.005 by whole-exome sequencing (case-control study) in the full epilepsy cohort (EPI). We next assessed whether expression of ASD-, NDD- and epilepsy-associated genes is regulated in *TSC2*-Null NPCs as determined by anota2seq analysis (i.e. from Figure 3A). This analysis revealed significant enrichments of ASD72 (∼43% of all ASD72 genes; p = 1.72×10^-7^) and NDD373 (∼37% of all NDD373 genes; p = 2.59×10^-19^) genes among translationally suppressed transcripts as compared to background (Table 1). This finding was further substantiated by applying two approaches that do not use thresholds to identify sets of differentially regulated genes: i) gene set enrichment analysis (GSEA) ^41^ (Figure 4A) and ii) the same method as applied for *TSC1* vs *TSC2* (i.e. Figure 3D) gene subset-comparisons (Figure 4B, top). Both these approaches showed strong enrichment of ASD72 and NDD373 genes among transcripts translationally suppressed in *TSC2*-Null as compared to isogenic WT NPCs. Similarly, in *TSC2*-Null NPCs, strong enrichments among translationally suppressed genes were observed for the larger ASD185 and NDD664 gene sets (Table S2A; Figure S3A, B). ASD- and NDD-associated genes were also enriched among transcripts that were translationally suppressed in both *TSC2*-Null and *TSC1*-Null NPCs relative to their respective controls (i.e. genes from Figure 3E; Table S2B). A similar analysis for EPI-associated genes revealed a smaller enrichment (∼27% of all EPI-associated genes; p = 5.82×10^-3^) (Table 1) that was not confirmed using methods that do not use thresholds to identify regulated sets of genes (Figure S3C, D).

**Table 1.**
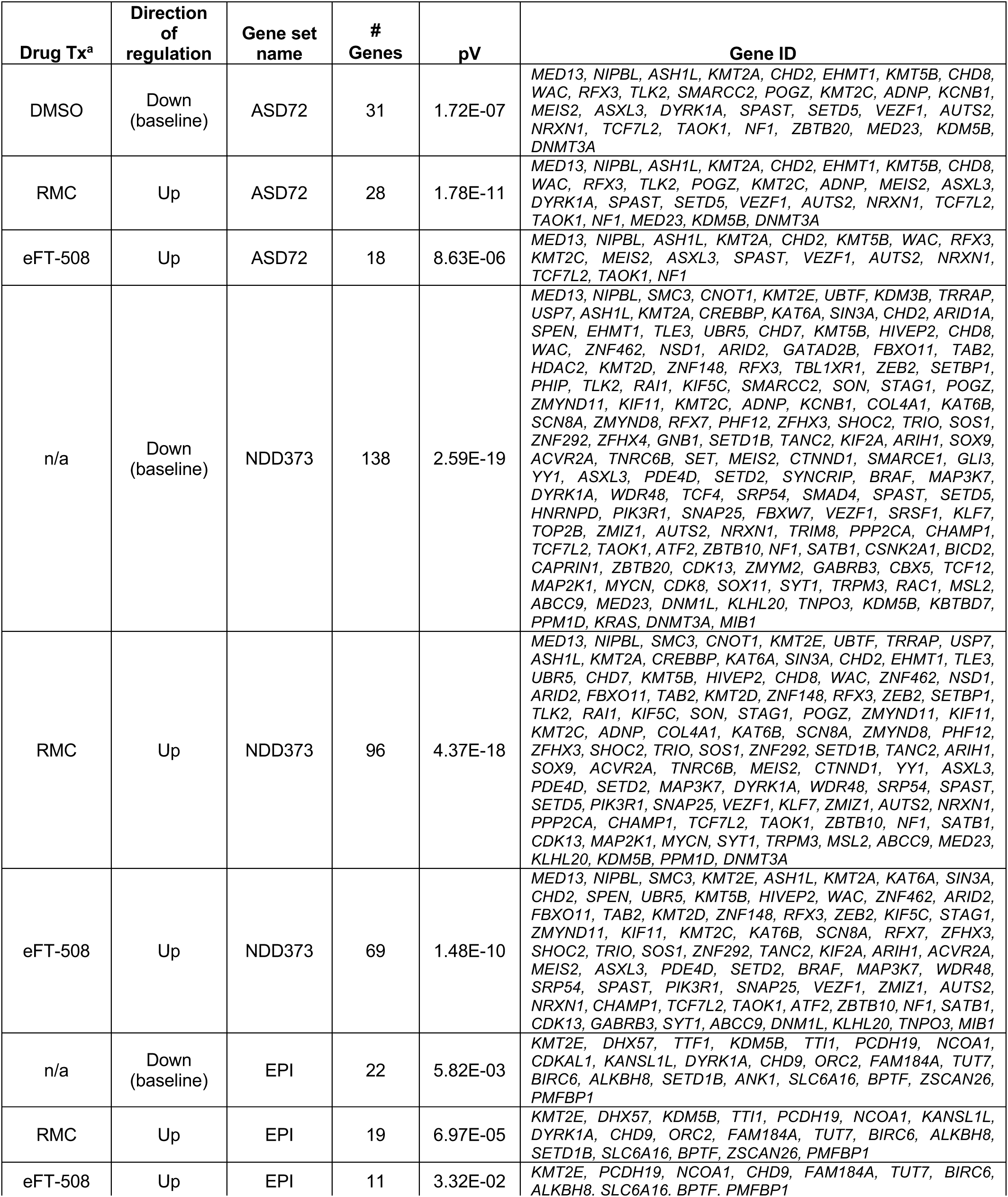
Gene set enrichment analysis of translationally downregulated genes in *TSC2*-Null NPCs related to ASD, NDD and epilepsy. ^a^Drug Tx, treatment: DMSO, 10 nM RMC-6272 or 50 nM eFT-508; n/a, not applicable; pV, p-value; ASD72, 72 gene variants (false discovery rate (FDR)<0.001) identified in patients with autism spectrum disorder; NDD373, 373 gene variants (FDR<0.001) identified in patients with neurodevelopmental disorders ^38^; EPI83, 83 gene variants identified in full epilepsy cohort ^42^.

**Figure 4.**
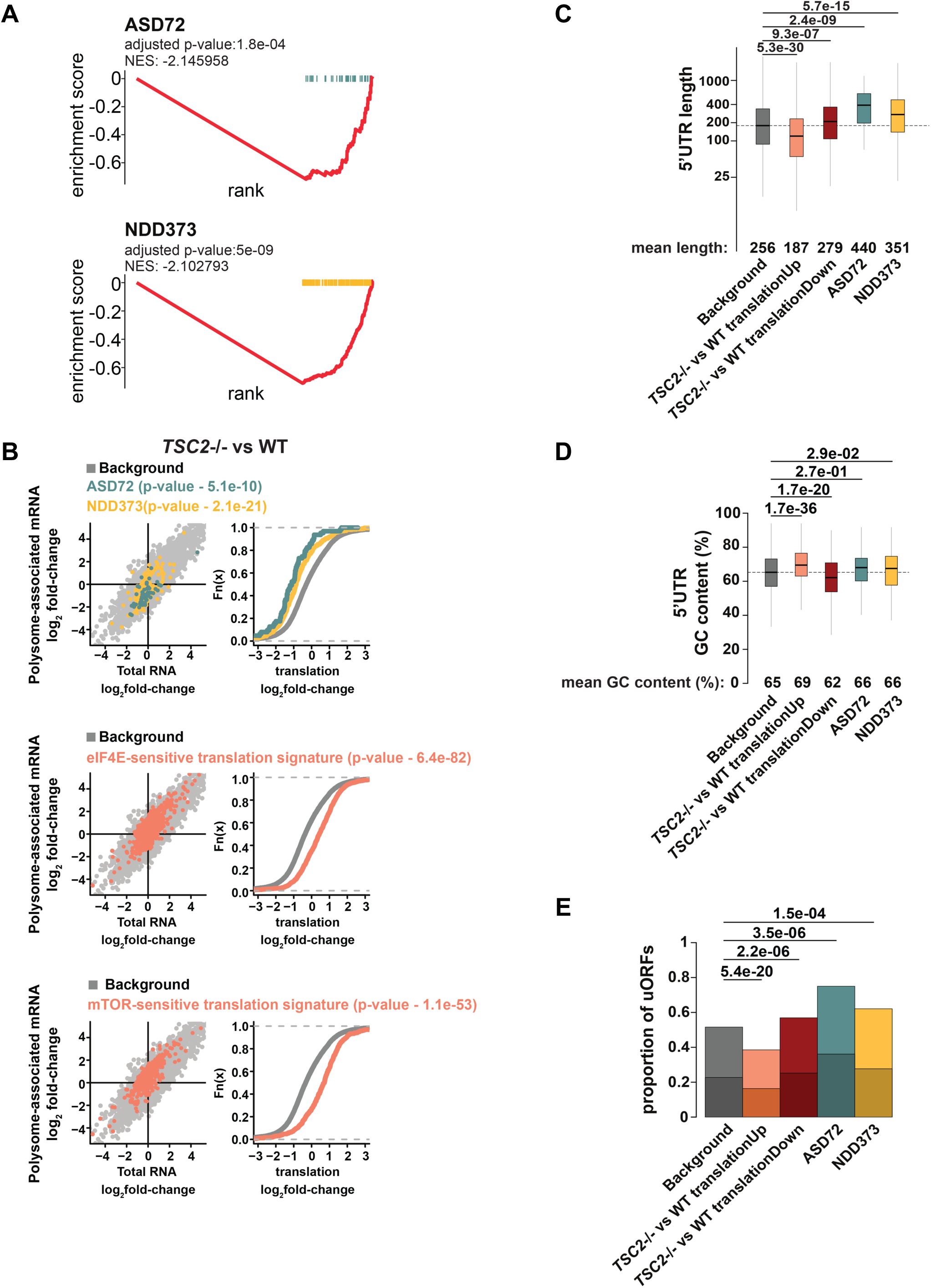
Suppressed translation of transcripts encoding ASD- and NDD-associated proteins in *TSC2*-Null NPCs associates with distinct 5’UTR features. (A) GSEA analysis of ASD- and NDD-associated genes using anota2seq analysis of translation (fold-changes) contrasting *TSC2*-Null (-/-) vs WT NPCs. Rankings for ASD- and NDD-associated genes (ASD72 and NDD373, respectively) were shifted towards the right denoting suppressed translation. Normalized enrichment scores (NES) and adjusted p-values are shown. (B) Scatterplots from anota2seq analysis for *TSC2*-Null vs WT NPCs. High confidence ASD- and NDD-associated genes (ASD72 and NDD373, respectively) (top left panel) as well as eIF4E- (middle left panel) or mTOR-sensitive (bottom left panel) transcripts are indicated. Shown to the right of each panel are corresponding empirical distribution functions together with associated statistics (as in Figure 3D). (C-D) Boxplots of 5’UTR lengths (C) and GC-contents (D) (according to the RefSeq database, version 109.20201120) for ASD- and NDD-associated genes as well as mRNAs translationally regulated in *TSC2*-Null vs WT NPCs (after excluding ASD- and NDD-associated genes; background corresponds to mRNAs absent in all subsets). The mean length or GC-contents for each set are indicated together with p-values from Wilcoxon rank-sum tests. (E) The proportion of mRNAs harboring upstream open reading frames (uORFs) in their 5’UTRs among subsets of transcripts from C-D. The lower part of the bar graphs corresponds to uORFs with a strong Kozak context. Differences in proportions of uORFs were assessed using Fisher’s exact tests for the indicated comparisons.

To assess whether this unexpected translational suppression of ASD- and NDD-associated genes in NPCs occurs in the presence of expected effects on mRNA translation from mTORC1 activation, we applied the same approaches described above to sets of transcripts whose translation was altered following overexpression of eIF4E ^39^. Consistent with activation of the mTORC1-eIF4F axis controlling translation in *TSC2*-Null NPCs, there was a strong overlap between genes whose translation was activated upon eIF4E-overexpression and *TSC2* loss (Table S3). A similar result was obtained when only considering genes that were identified as translationally regulated by anota2seq following both *TSC1* and *TSC2* loss (i.e. from Figure 3E; Table S3) and when a method not relying on thresholds was used (Figure 4B, middle) as well as when using an additional signature of mTORC1-sensitive translation reported previously ^14^ (Figure 4B, bottom). Therefore, interestingly, genes associated with ASD and NDD are largely translationally suppressed in NPCs following inactivation of *TSC2* despite appropriate activation of mTORC1-sensitive translation.

### 5’UTRs of ASD and NDD genes resemble those of mRNAs whose translation is suppressed upon activation of mTORC1

The above results unveiled that translation of mRNAs encoding proteins associated with ASD and NDD is affected by the activity of mTORC1. As mTORC1-sensitive translation largely relies on features in the 5’UTR, this suggests that the observed suppressed translation of ASD- and NDD-associated genes may depend on their 5’UTRs. As mentioned above, mRNAs whose translation parallels activity of mTORC1 often have short and/or highly structured 5’UTRs and lack upstream open reading frames (uORFs). Strikingly, ASD- and NDD-associated genes showed an opposite profile for 5’UTR length, GC-content (which associates with secondary structures of the 5’UTR) and uORFs that was more similar to mRNAs whose translation was suppressed upon *TSC2* loss (Figure 4C-E). Therefore, ASD- and NDD-associated genes largely show 5’UTRs that are consistent with translational suppression following increased activation of mTORC1.

### RMC-6272 and eFT-508 treatment reverses *TSC2*-dependent translation and encompasses ASD- and NDD-associated genes

The findings above suggest that suppressed translation of ASD- and NDD-associated genes following loss of *TSC2* may be important for neurodevelopmental manifestations. Furthermore, this raises the possibility that inhibition of mTORC1 signaling, or potentially other key pathways modulating translation, may reverse *TSC2*-dependent translation of ASD- and NDD-associated genes. To assess this, we performed polysome-profiling in *TSC2*-Null NPCs treated with mTORC1 inhibitor RMC-6272, or the MNK1/2 inhibitor eFT-508. The resulting dataset was analyzed as described above using anota2seq to determine how these agents modulate gene expression. Both treatments resulted in dramatic mRNA translation changes as visualized in scatterplots comparing effects in polysome-associated and total mRNA levels as well as p-value and FDR densities (Figure S4A-D). Next, we generated gene signatures of transcripts that were translationally increased or decreased in *TSC2*-Null NPCs treated with RMC-6272 relative to DMSO control and determined how these gene sets were regulated in *TSC2*-Null NPCs treated with eFT-508 vs DMSO control. This revealed that transcripts whose translation was activated or suppressed by RMC-6272 showed similar regulation upon eFT-508 treatment and, vice versa, transcripts whose translation was activated or suppressed by eFT-508 showed similar regulation upon RMC-6272 treatment (Figure 5A, B). Yet, these overlaps were partial as subsets of transcripts that were altered in one treatment group did not shift in the other treatment group (Fig. 5A and B). To determine whether the agents reversed *TSC2-*sensitive translation, we used the same strategy as above when comparing *TSC1-* vs *TSC2-*sensitive translation (i.e. Figure 3D). This revealed that both RMC-6272 as well as eFT-508 partially reversed *TSC2-*sensitive translation (Figure 5C). Gene ontology analysis of genes overlapping in *TSC2*-Null NPCs treated with RMC-6272 or eFT-508 revealed significant reversal of some cellular functions that were identified as *TSC2*-dependent (shown in Figure 3C), including positive regulation of transcription by RNA polymerase II, positive regulation of RNA metabolic process and protein DNA complex organization along with transcripts involved in various metabolic processes (Figure 5D).

**Figure 5.**
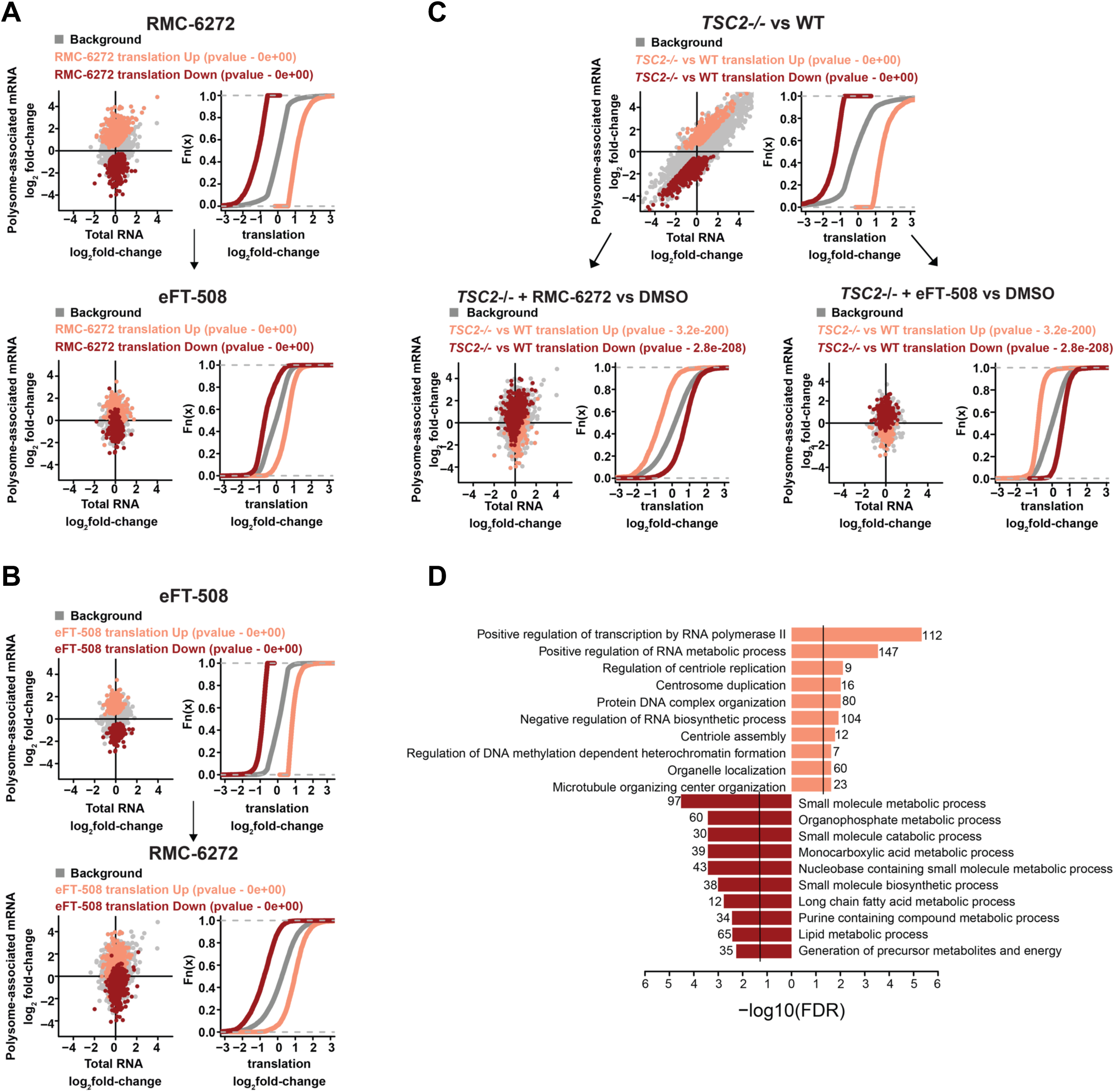
mTORC1 or MNK1/2 inhibition reverses the TSC2-associated translatome. (A and B) Scatterplots from anota2seq analysis of *TSC2*-Null (-/-) NPCs treated with RMC-6272 vs DMSO (A, top left panel) or eFT-508 vs DMSO (B, top left panel). Sets of transcripts whose translation was activated or suppressed under each treatment were then assess across treatments (A and B, bottom panels). Also shown are empirical cumulative distribution functions (A and B, right panels) of the log2 fold changes in translation for the corresponding gene sets. Grey dots and lines correspond to the background (i.e. absent in gene sets). Significant shifts of gene sets, relative to background, were determined using the Wilcoxon rank-sum test and the resulting p-values are shown. (C) Assessment of how transcripts whose translation was altered in *TSC2*-Null vs WT NPCs (top panel, from Figure 3A) are modulated upon treatment with RMC-6272 (bottom left panel) or eFT-508 (bottom right panel). (D) Enrichments of gene ontology annotation for biological processes among genes identified with reversed translation, relative to baseline (*TSC2*-Null vs WT NPCs) that overlap between RMC-6272 and eFT-508 treatment (same color codes as in Figure 3C).

By comparing anota2seq outputs for these analyses, we observed that among all transcripts whose translation was increased in *TSC2*-Null cells (Figure 3A), 614 (∼47%) or 936 (∼71%) were suppressed upon RMC-6272 or eFT-508 treatment, respectively, with an overlap of 513 transcripts. Similarly, among all transcripts whose translation was decreased in *TSC2*-Null cells, 1454 (∼57%) or 1177 (∼46%) were translationally activated upon RMC-6272 or eFT-508 treatment, respectively, with an overlap of 874 transcripts (all overlaps were statistically significant; Figure S4E; Table S4). Gene ontology analyses of *TSC2*-associated changes in translation (from Fig. 3A) that were distinctly reversed upon treatment with either RMC-6272 or eFT-508 revealed significant reversal of enriched annotations upon treatment with RMC-6272 or eFT-508 as well as reversal of key cellular functions shown in Figure 3C (Figure S4F, G).

Although the inhibitors led to altered translation of similar numbers of transcripts in *TSC2*-Null NPCs, anota2seq analysis contrasting the two treatments revealed notable differences in terms of affected mRNAs (Figure S5A, B). This was further substantiated by gene ontology analyses, which revealed that while RMC-6272 more efficiently suppressed translation of mRNAs encoding proteins involved in translation and oxidative phosphorylation, eFT-508 more efficiently suppressed translation of mRNAs relating to neuronal development (Figure S5C).

Next, we assessed whether translation of mRNAs encoding ASD- and NDD-associated genes were affected by mTORC1 and/or MNK1/2 inhibition. This revealed a striking reversal in translation of both ASD and NDD genes in *TSC2*-Null NPCs treated with RMC-6272 or eFT-508 (Figure 6A). To identify affected genes, we compared outputs from the corresponding anota2seq analyses. Among the 31 ASD72 genes translationally suppressed in *TSC2*-Null NPCs, 28 (∼90%) and 18 (∼58%) were translationally activated upon RMC-6272 (RMC-responsive) and eFT-508 treatment (eFT-responsive), respectively. While all eFT-responsive genes were identified as RMC-responsive, 10 genes were only responsive to RMC-6272 (Table 1; Figure 6B). Moreover, for translationally suppressed NDD373 genes, 96 (∼70%) and 69 (50%) were RMC- and eFT-responsive (Table 1), respectively (60 were both RMC- and eFT-responsive; similar results were obtained for ASD185 and NDD664 genes; Table S2A). To validate alterations in translation upon *TSC2* loss as well as the reversal following mTORC1 or MNK1/2 inhibition, we selected five representative ASD/NDD-associated genes that were translationally suppressed in *TSC2*-Null NPCs: *SETD1B* [MIM: 611055], *DYRK1A* [MIM: 600855], *CHD8* [MIM: 610528], *NRXN1* [MIM: 600565] and *TCF7L2* [MIM: 62228], and also five eIF4E-sensitive genes that were translationally activated in *TSC2*-Null NPCs: *AP2S1* [MIM: 602242], *LZTR1* [MIM: 600574], *WDR54*, *ALDOC* [MIM: 103870] and *EIF2B2* [MIM: 606454]. Quantitative RT-PCR (RT-qPCR) analysis from polysome-associated mRNA fractions (>2 ribosomes), adjusted for alterations in total mRNA, normalized to housekeeping gene, *ACTB* showed a reduction for all five ASD/NDD-related genes and increase in all five eIF4E-sensitive genes in *TSC2*-Null vs WT NPCs, confirming the above polysome-profiling results. Moreover, RMC-6272 or eFT-508 reversed *TSC2*-associated translationally suppressed ASD/NDD and translationally activated eIF4E-sensitive genes (Figure 6C). Similar results were obtained when normalized to a second housekeeping gene *PGK1* (data not shown).

**Figure 6.**
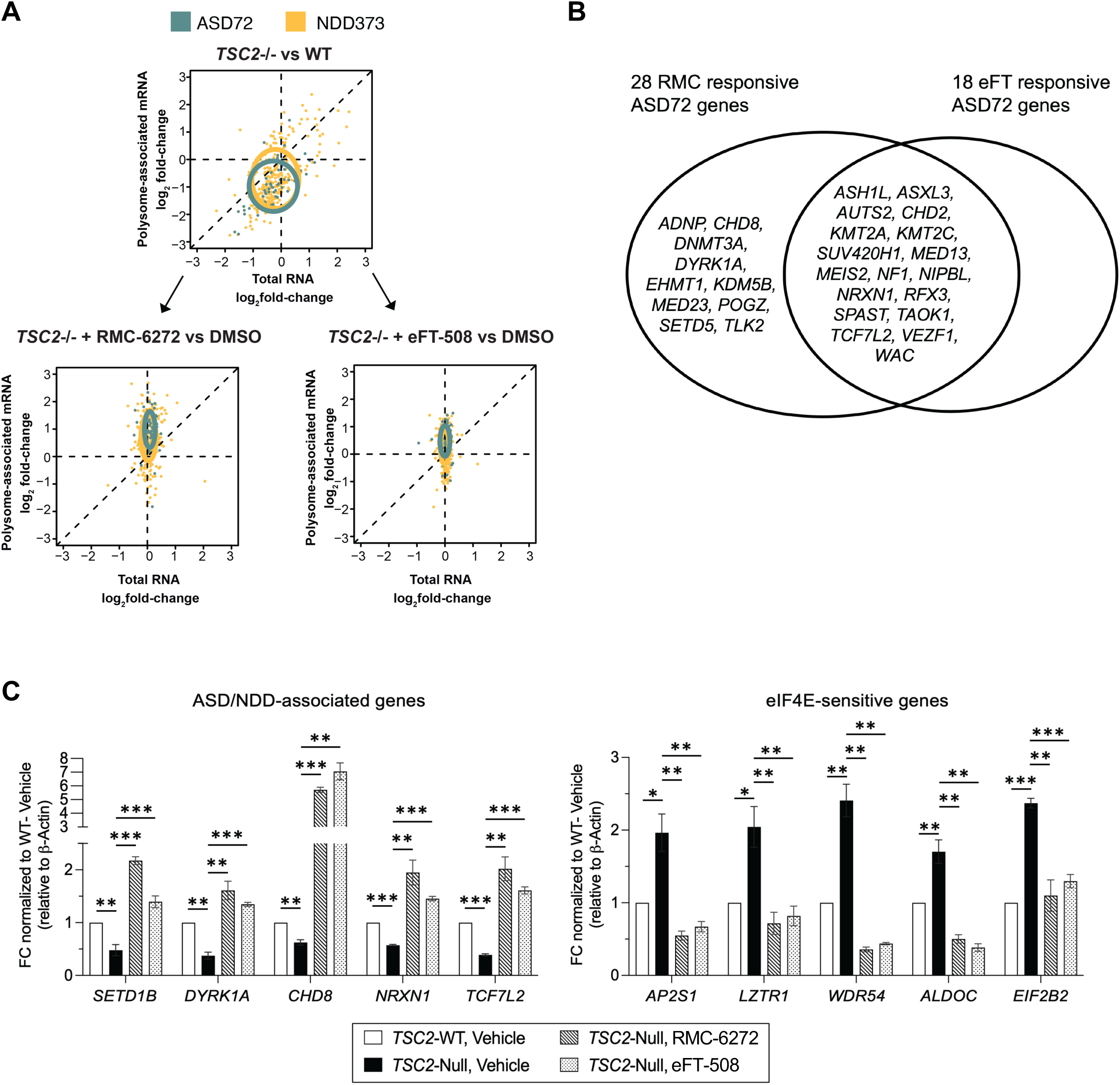
mTORC1 or MNK1/2 inhibition reverses translation of transcripts encoding ASD- and NDD-associated proteins in *TSC2*-Null NPCs. (A) Scatterplots, showing high confidence ASD- and NDD-associated genes (ASD72 and NDD373), from anota2seq analysis comparing *TSC2*-Null (-/-) vs WT NPCs (top panel) as well as RMC-6272 (bottom left) or eFT-508 (bottom right) treated relative to DMSO treated *TSC2-* Null NPCs are shown. The shape and position of ellipses correspond to the mean and standard deviation, respectively, for all genes within each gene set. (B) A Venn diagram of ASD associated genes (ASD72) whose translation was suppressed in *TSC2*-Null NPC relative to *TSC2*-WT NPCs and reversed by treatment with RMC-6272 or eFT-508. (C) Plots denote RT-qPCR analysis from the polysome-associated RNA pool (3 biological replicates) for 5 ASD- or NDD-associated genes (left) as well as 5 eIF4E-sensitive genes (right) from the indicated cells and treatments. FC, fold change. Data were normalized to total mRNA obtained in parallel as well as ý-actin *(ACTB)* mRNA expression. P-values from student’s t-test test (*p<0.05, **p<0.01, ***p<0.001) for the indicated comparisons are shown.

Taken together, *TSC2*-dependent decreased translation of mRNAs associated with ASD and NDD can be reversed upon RMC-6272 or eFT-508 treatment in NPCs. These results are intriguing and potentially imply paradoxically suppressed translation of a subset of neurodevelopmentally important genes that nonetheless depend on mTORC1-eIF4F pathway activation.

## DISCUSSION

The central role of mTORC1 signaling in translation initiation and protein synthesis in a variety of cell types including stem cells, as well as in human disease, is well established ^11; 50;51^. While many transcriptome-wide studies have been reported in *TSC1/2*-deficient neuronal cells, how *TSC2* loss affects mRNA translation, particularly in patient-derived cells, is unknown. We recently reported *TSC1*-dependent alterations in mRNA translation employing an isogenic set of CRISPR-modified, TSC1 patient-derived NPCs. Although polysome-profiling revealed a partial reversal of *TSC1*-associated gene expression changes following rapamycin treatment, most genes related to neural activity/synaptic regulation or ASD remained rapamycin-insensitive ^25^. Here, we describe generation and characterization of an independent isogenic set of CRISPR-modified NPCs from a TSC patient with a loss-of-function germline mutation in *TSC2*. We find that *TSC2*-Null NPCs mimic *TSC1* loss in activation of mTORC1 signaling and early neurodevelopmental phenotypic changes ^25^. Rapamycin treatment of *TSC2*-Null NPCs fully inhibited phosphorylation of S6K and reduced enlarged cell size but had minimal effects on 4E-BP1 phosphorylation and early neurodevelopmental phenotypes. In contrast, the third-generation mTORC1 inhibitor RMC-6272 revealed efficient inhibition of both S6K and 4E-BP1 phosphorylation and rescued these phenotypes as well as cell size. These data suggest that while enlarged cell size observed upon *TSC2* loss depends on the mTORC1-S6K axis of the pathway, which is consistent with previous reports ^47^, the altered neurodevelopmental phenotypes depend on the mTORC1-4E-BP1 signaling axis.

We observed dramatic changes in mRNA translation associated with *TSC2* loss in NPCs that largely overlapped with those observed upon *TSC1* loss. Provided that *TSC1* and *TSC2* loss assert similar effects on mTORC1 signaling, a substantial overlap is expected. Importantly as the cells were derived from independent patients, our results support a robust effect on mRNA translation across patients with hyperactive mTORC1 activity. In this study, in addition to RMC-6272, which inhibits mTORC1 activity, we also considered alternative approaches to modulate activity of the eIF4F complex and therefore included the MNK1/2 inhibitor eFT-508/tomivosertib to understand a possible convergence of mTORC1-4E-BP1 and MNK-eIF4E signaling pathways on *TSC2*-dependent protein synthesis (see Figure 1A). While mTORC1-sensitive translation has been studied extensively at the transcriptome-wide level ^22; 40; 52; 53^, the understanding of how the MNK-eIF4E axis modulates mRNA translation at a global scale is more limited ^16^. This study is the first that includes a direct comparison of these pathways to assess effects on mRNA translation at a transcriptome-wide level, and the results reveal that a large proportion of translational alterations observed in *TSC2*-Null NPCs can be reversed by RMC-6272 or eFT-508, suggesting that these changes are mediated by mTORC1-4E-BP1- and MNK-eIF4E-dependent activation of eIF4F complex. However, the activity of the two inhibitors is not equivalent since RMC-6272 would be expected to affect both eIF4E-dependent and S6K-dependent translation whereas eFT-508 would be restricted to MNK1/2-dependent translation. We also observe ample differences between the two drugs with respect to translation, which are in line with previous studies showing differences in affected cellular phenotypes as the mTORC1-4E-BP1 axis is a key regulator of cell cycle progression ^19; 54^ while MNK1/2-eIF4E controls cell migration ^55; 56^.

As epilepsy and ASD are strongly associated with TSC, we compared transcripts that were differentially translated in *TSC2*-Null vs WT NPCs to recent genetic datasets available from large cohorts of individuals with ASD, NDD and epilepsy ^38; 42^. Interestingly, numerous non-monogenic ASD-, NDD- and epilepsy-associated genes identified in patients harboring putative loss-of-function mutations, including protein truncating or damaging missense variants, were translationally suppressed in *TSC2*-Null NPCs and overlapped with data from our previously reported *TSC1*-model. Notably, the results for ASD- and NDD-associated genes were confirmed using multiple methods, which suggests that for these subsets there is a strong negative regulatory effect on mRNA translation impacting a large proportion of these mRNAs. This was not the case for epilepsy-associated genes, in that a smaller fraction of these genes was translationally suppressed upon *TSC2* loss. Importantly, these decreases in translation were observed side-by-side with the expected increased translation of mRNAs that were previously reported to be translationally activated upon mTORC1 pathway activation. This suppressed translation was, for a majority of identified ASD/NDD genes, reversed by inhibition of mTORC1 using RMC-6272 or MNK1/2 using eFT-508. A recent elegant large-scale transcriptome study demonstrated that CRISPR perturbation of several ASD genes results in differential expression of ASD/NDD-related genes that positively correlate with co-expression of the perturbed gene, thereby establishing a high level of convergence across ASD risk genes ^57^. Our results here showing translational suppression of many ASD/NDD genes upon loss of *TSC2* is consistent with the positive correlation and co-expression reported by Liao and colleagues ^57^. However, most notably, our results demonstrate that a majority of translationally suppressed ASD/NDD/epilepsy genes can be reversed by RMC-6272 or eFT-508, thereby establishing the roles of mTORC1-4E-BP1 and MNK-eIF4E in regulating translation of these genes.

Many of the translationally suppressed ASD genes are FMRP targets. FMRP was originally considered to function as a translational repressor that can bind to numerous mRNAs that are generally larger than average and enriched for genes associated with ID and ASD ^58;59^. However, recent studies of FMRP-deficiency have documented a decrease in translation initiation rates of many FMRP targets including ASD-related genes, which contrasts the notion that FMRP is solely a translational repressor and supports a model where FMRP can also function to increase the translation initiation of many targets ^60–63^. Furthermore, in *TSC2*-Null NPCs we also observe that 32 translationally suppressed ASD/NDD-related genes are associated with chromatin-remodeling ^64; 65^. Additional studies are therefore necessary to fully comprehend how the ASD- and NDD-associated genes are translationally suppressed upon loss of *TSC2*.

In the literature, the focus has been, in general, towards understanding mechanisms underlying transcript-selective translation paralleling mTORC1 activity ^13^ while less attention has been given to transcript that show decreased translation upon mTORC1 activation. Translation of mRNAs enriched for 5’UTRs containing uORFs are translationally suppressed upon mTORC1 activation ^40^. In agreement, we observed an increased frequency of uORFs in both ASD- and NDD-associated genes; however, it should be noted that uORFs are also common among background transcripts, and a relatively large proportion of ASD- and NDD-associated genes do not have uORFs in their 5’UTRs.

Our results raise the critical question as to how ASD/NDD-related genes are translationally suppressed upon mTORC1 activation in *TSC2*-Null NPCs. In addition to mTORC1 signaling, the integrated stress response (ISR) pathway, which selectively modulates translation of transcripts with 5’UTRs harboring uORFs ^66^, is a key regulator of mRNA translation. In response to cellular stress, a decrease in global protein synthesis is paralleled by increased translation of mRNAs with 5’-uORFs including the transcription factor ATF4 (reviewed in ^67^); therefore, further studies are need to examine involvement of the ISR pathway in our TSC models. Furthermore, alternative mechanisms such as DAP5-dependent translation, which can be activated under low mTORC1 signaling, may also play a role ^53^.

Another potential mechanism of translational regulation is via microRNAs (miRNAs), which are involved in neurodevelopmental disorders ^68^. Among 31 translationally suppressed transcripts in *TSC2*-Null NPCs that show strong association with ASD (ASD72, FDR<0.001), 28 are predicted to be targeted by at least one miRNA, using the miTarBase 2017 library (data not shown). Interestingly, of the individual miRNAs identified by miTarBase analysis, four (hsa-miR-27b-3p, hsa-let-7g-3p, hsa-miR-29a-3p and hsa-miR-515-5p) were reported as increased upon *TSC1*- or *TSC2*-shRNA suppression in human neural stem cells generated from skin fibroblast-derived iPSCs (p <0.05) ^69^; and three (hsa-miR-21-5p, hsa-miR-19b-3p and hsa-miR-144-3p) were shown to be overexpressed in human autism brain samples ^70^. Based on this, it is tempting to speculate that miRNAs may play a role in suppressing translation in *TSC2*-Null NPCs, and future studies will address this mechanistic possibility.

In summary, we establish that loss of *TSC2* in patient-derived NPCs results in numerous changes in mRNA translation, which significantly overlap with loss of *TSC1*. Loss of *TSC2* associates with suppressed translation of a large proportion of ASD- and NDD-related genes, which is reversed by inhibition of mTORC1 or MNK1/2, demonstrating the importance of eIF4F-dependent mRNA translation in neurodevelopmental disorders. Our work also supports the therapeutic potential for RMC-5552 (the clinical counterpart of RMC-6272) and eFT-508/tomivosertib that are currently under clinical development for cancer, as a strategy for treatment of TSC-associated neuropsychiatric disorders as well as ASD and other neurodevelopmental disorders.

## ACKNOWLEDGEMENTS

This paper is dedicated to the memory of the late Dr. Jerry Pelletier, a distinguished James McGill Professor in the Department of Biochemistry and Oncology from the Goodman Cancer Institute at McGill University, Montreal, Canada, who unfortunately passed away during the completion of this study. This work was supported by NIH R01 NS109540 (V.R.), the Swedish Brain foundation FO2022-0178 (O.L.); the Swedish research council 2020-01665 (O.L.); the Wallenberg academy fellow program 2013.0181 (O.L.). We thank Revolution Medicines, Inc. for generously providing RMC-6272 for this study. Our sincere thanks to Dr. Justin Meyerowitz and Dr. Mallika Singh of Revolution Medicines, Inc. for their valuable comments on the manuscript.

## AUTHOR CONTRIBUTIONS

Conceptualization and study design: V.R. and O.L.; experimental methods and data acquisition P.M., K.S., F.R., S.B., R.B., N.R., J.B., S.C. and I.N.; data analysis and interpretation: P.M., K.S., F.R., S.B., R.B. S.C., I.N., S.H., S.E. and O.L.; manuscript preparation and editing: P.M., K.S., F.R., S.B., R.B., I.N., S.H., S.E., O.L. and V.R.

## WEB RESOURCES

OMIM, https://www.omim.org/

NCBI GenBank, https://www.ncbi.nlm.nih.gov/genbank/

Sequences,https://www.ncbi.nlm.nih.gov/nuccore/NM_000548

GenScript, https://www.genscript.com

BBTools software suite, https://www.osti.gov/servlets/purl/1241166

fgsea package (v.1.24.0), https://doi.org/10.18129/B9.bioc.fgsea

R statistical software environment, https://www.R-project.org/

## DATA AND CODE AVAILABILITY

The polysome-profiling datasets supporting the conclusions of this article are included within the article (and its additional file(s)), and the full datasets will be deposited into Gene Expression Omnibus (GEO).

