## Supplemental Figures S1-S5 for "*TSC2* loss in neural progenitor cells suppresses translation of ASD/NDD-associated transcripts in an mTORC1- and MNK1/2-reversible fashion"

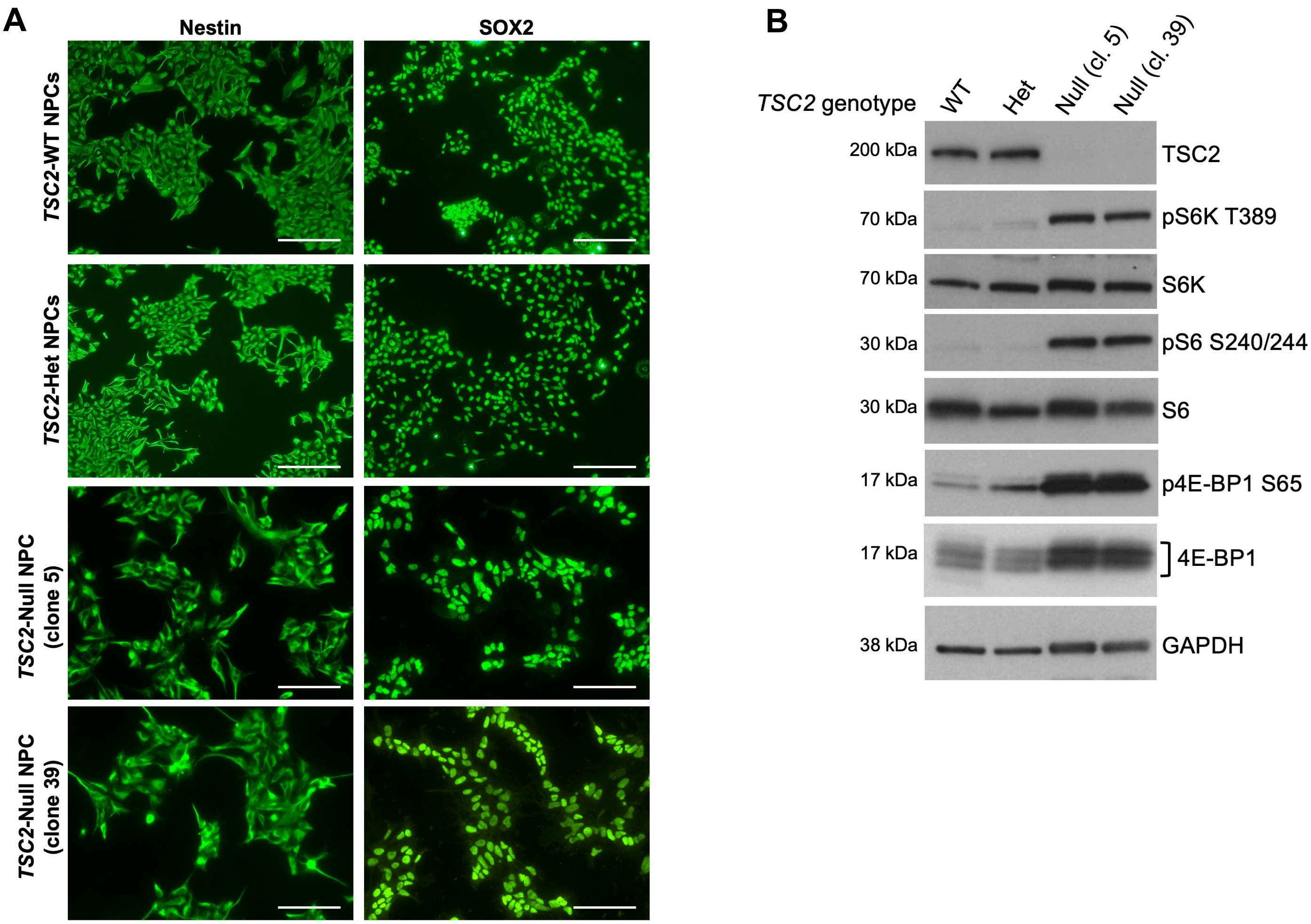


**Supplemental Figure S1. Characterization of TSC2 iPSC-derived NPCs. A.** An isogenic set of isogenic *TSC2*-NPCs (Het, Null, WT) generated from skin fibroblast-derived iPSCs, express neural progenitor markers Nestin (left panel) and SOX2 (right panel). Immunostaining was performed at least 3 times. Scale bar=100μm. **B.** Immunoblotting of TSC2-NPCs, including one clone each for *TSC2*-WT and *TSC2*-Het, and two independent clones (cl. 5 and 39) for *TSC2*-Null, is shown for TSC2/tuberin as well as mTORC1 pathway readouts pS6K T389, pS6 S240/244 and p4E-BP1 S65. GAPDH and respective total proteins serve as controls.

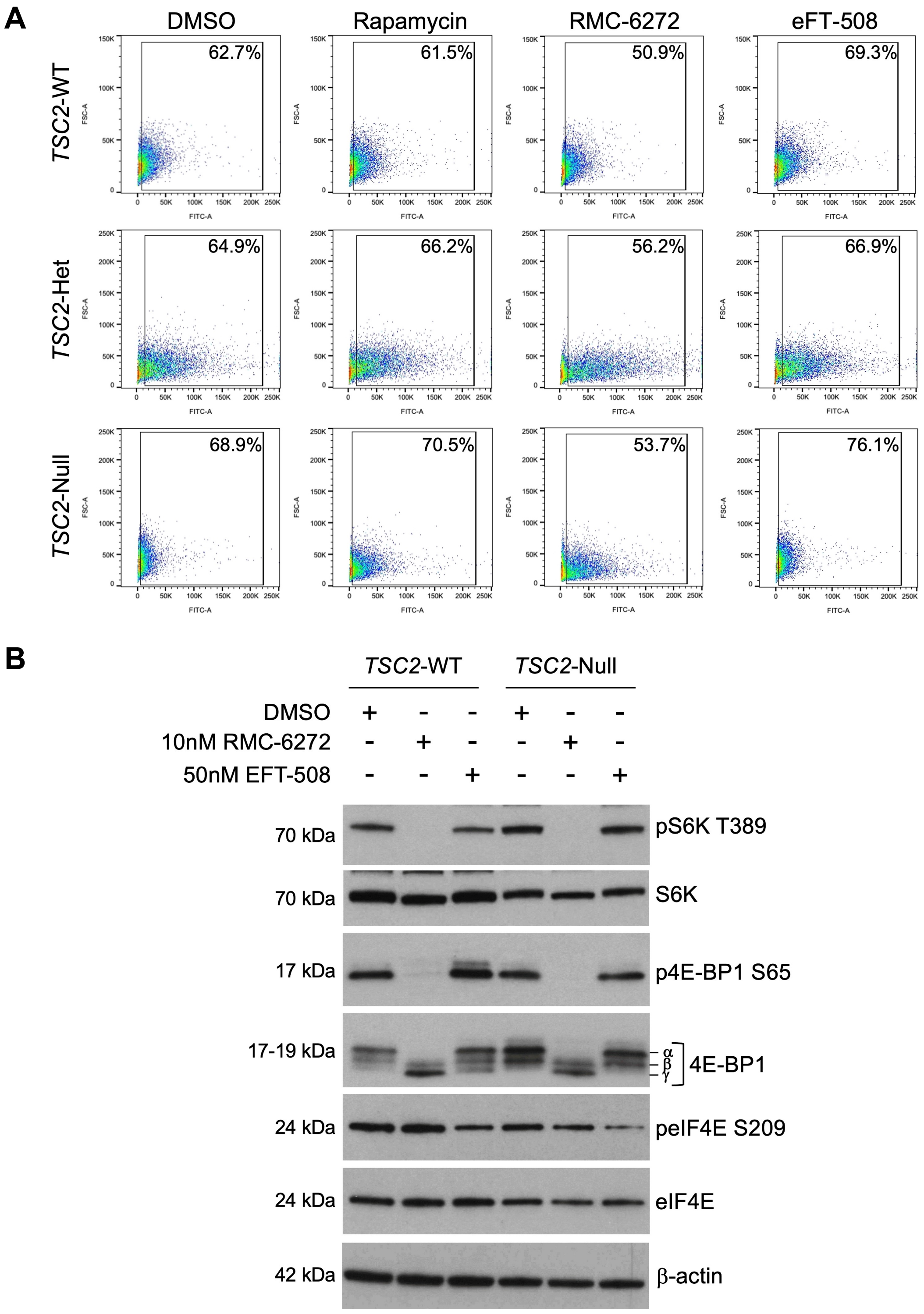


**Supplemental Figure S2. Treatment of TSC2-NPCs with RMC-6272 or eFT-508. A.** Representative flow cytometry plots for Ki-67-positive cell populations are shown for of *TSC2*-WT, -Het and -Null NPCs, are shown under control conditions (DMSO) or 72 h treatment with rapamycin (50 nM), RMC-6272 (10 nM) or eFT-508 (50 nM). Graphs represent plotting of forward scatter-area (FSC-A) versus FITC-area from two biological replicates, indicating positivity of cells for Ki-67-Alexa488. Boxed regions show gating of Ki-67-positive staining along with indicated percentages. **B.** Immunoblotting of *TSC2*-WT and *TSC2*-Null NPCs following 2 h treatment with DMSO (control), rapamycin (50nM) or RMC-6272 (10nM) is shown for mTORC1 targets pS6K T389 and p4E-BP1 S65 as well as MNK1/2 target peIF4E S209. β-actin and respective total proteins serve as controls.

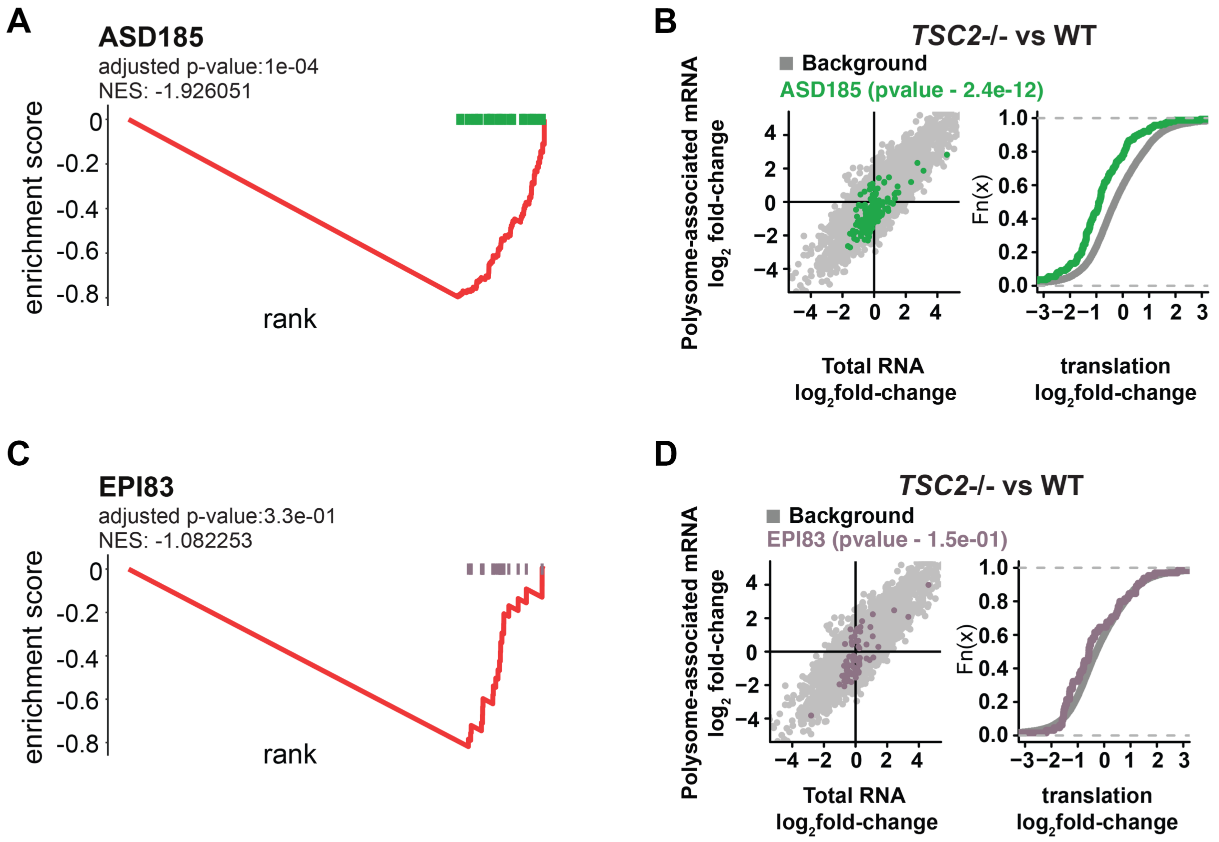


**Supplemental Figure S3. GSEA analysis of ASD- and Epilepsy-associated genes.**
**A-D.** GSEA analysis of ASD- (ASD185, A and B) and epilepsy- (EPI83, C and D) associated genes and anota2seq analyses of translation (fold-changes) from the comparison of *TSC2*-Null (-/-) versus WT NPCs. Rankings for ASD- and epilepsy-associated genes (ASD185 and EPI83, respectively) were shifted towards the right, indicating suppressed translation. Normalized enrichment scores (NES) and adjusted p-values are shown (A and C). The GSEA plots are accompanied by scatterplots from the anota2seq analysis for *TSC2*-Null versus WT NPCs. High-confidence ASD- and epilepsy-associated gene (ASD185 and EPI83, respectively) transcripts are indicated (B and D, left panels). Shown to the right of each panel are the corresponding empirical distribution functions along with associated statistics (as in Fig. 3D).

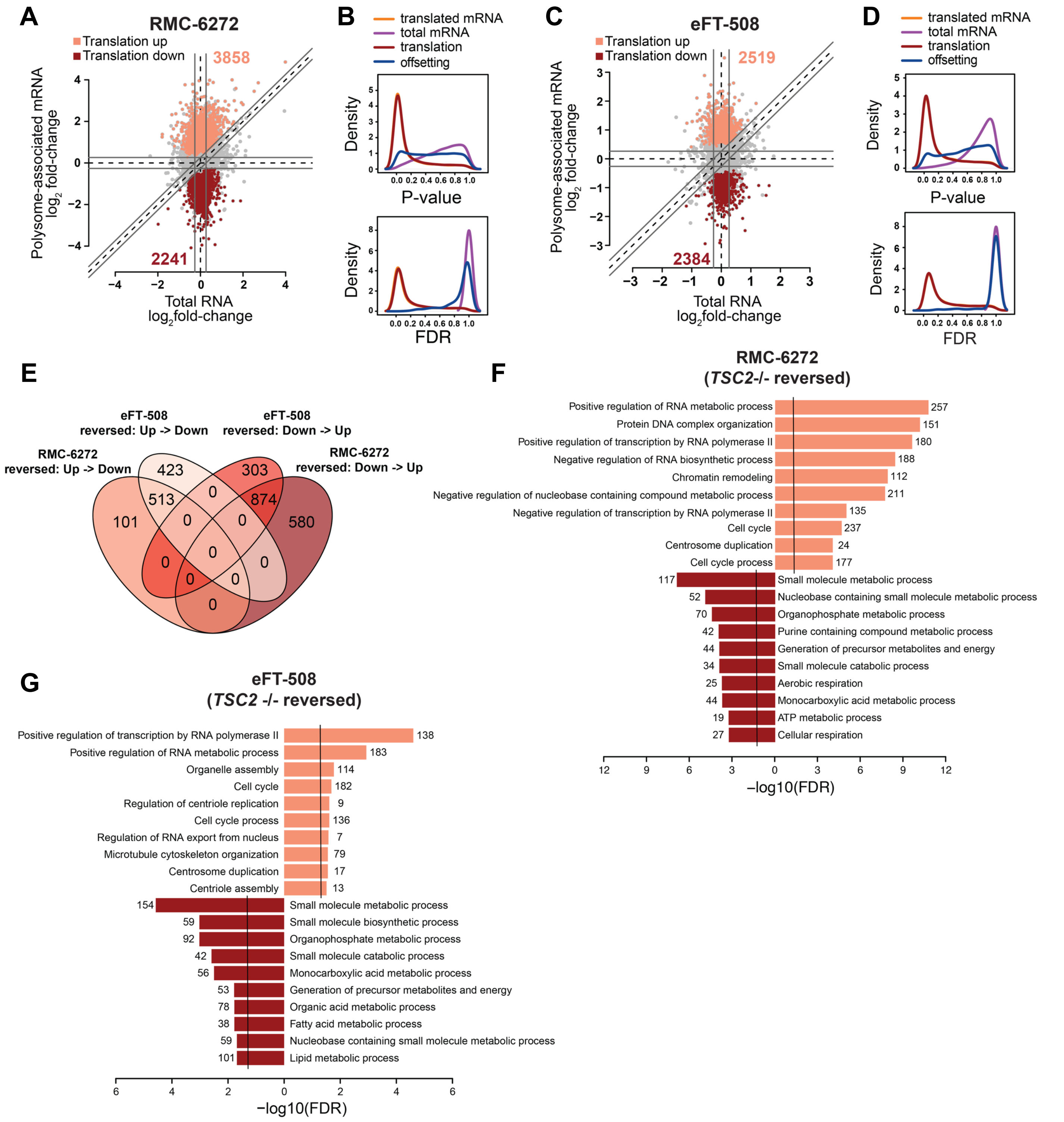


**Supplemental Figure S4. Effect of Treatment with RMC-6272 or eFT-508 on mRNA Translation in *TSC2*-Null NPCs. A-D.** Scatter plots (A and C) of polysome-associated mRNA versus total mRNA log2 fold changes between *TSC2*-Null NPCs treated with RMC-6272 versus DMSO (A) and eFT-508 versus DMSO (C). Genes are colored by mode of regulation as identified by the anota2seq analysis (FDR < 0.15). The number of mRNAs regulated via translation is indicated. Kernel densities of p-values and FDRs (from anota2seq analysis, B and D) for respective comparisons of *TSC2*-Null NPCs treated with RMC-6272 or eFT-508. Densities are shown for the analysis of polysome-associated RNA (translated mRNA), total mRNA, translation, and offsetting. A shift of the density towards low p-values/FDRs indicates a higher frequency of changes. **E.** Venn diagram comparing number of genes whose translation was altered in *TSC2*-Null relative to WT NPCs and reversed by treatment with RMC-6272 or eFT-508. **F and G.** Enrichment of gene ontology annotations for biological processes among genes identified as showing reversal in translation RMC-6272 (F) or eFT-508 (G) treatment.

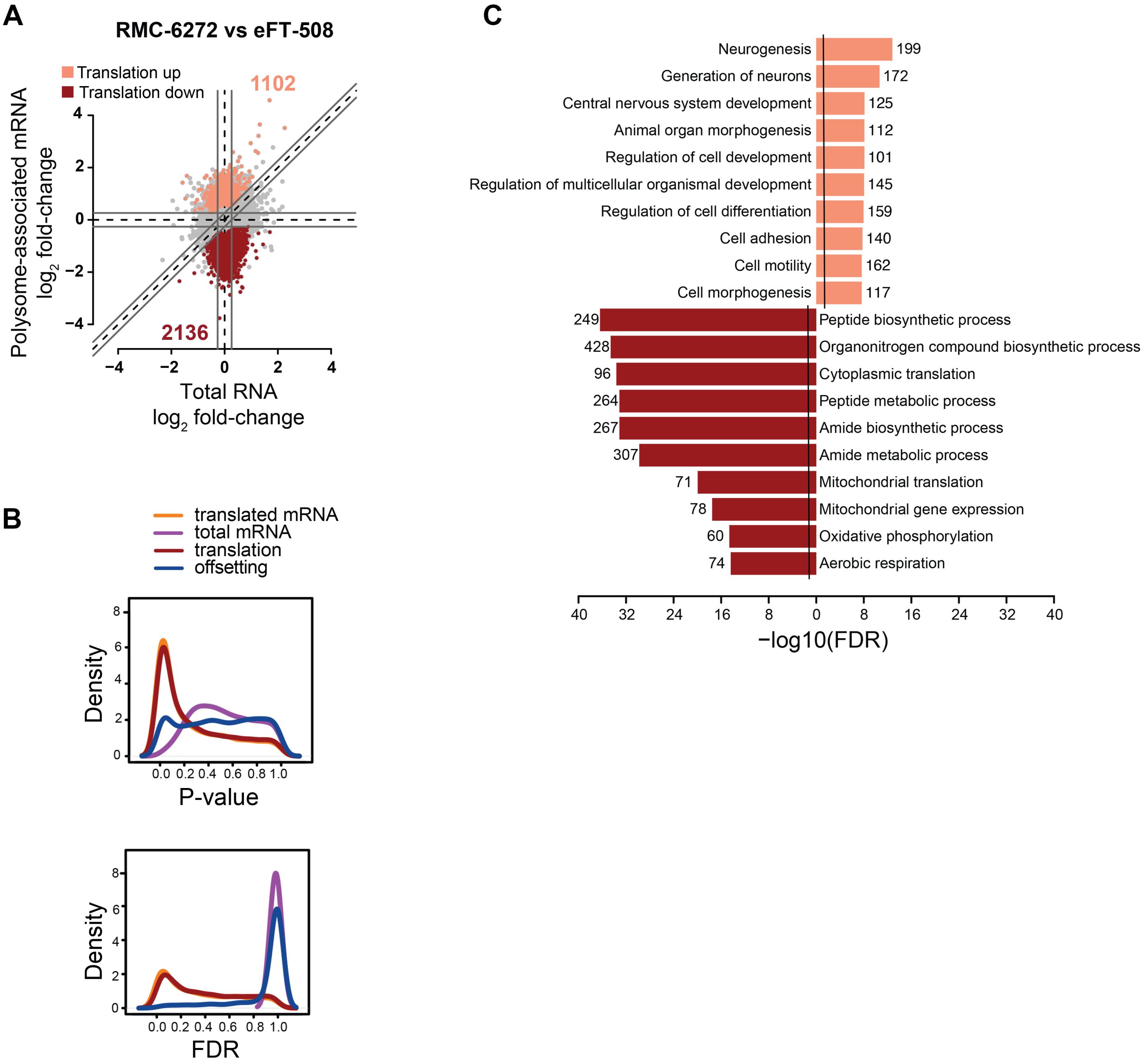


**Supplemental Figure S5. Comparison of Differences between Treatment with RMC-6272 and eFT-508 on mRNA Translation in *TSC2*-Null NPCs. A.** Scatter plots of polysome-associated mRNA versus total mRNA log2 fold changes between *TSC2*-Null NPCs treated with RMC-6272 versus eFT-508. Genes are colored according to their mode of regulation as identified by the anota2seq analysis (FDR < 0.15). The number of mRNAs regulated via translation is indicated. **B.** Kernel densities of p-values and FDRs (from anota2seq analysis) for the comparison of *TSC2*-Null NPCs treated with RMC-6272 versus eFT-508. Densities are shown for the analysis of polysome-associated RNA (translated mRNA), total mRNA, translation, and offsetting. A shift of the density towards low p-values/FDRs indicates a higher frequency of changes. **C.** Enrichment of gene ontology annotations for biological processes among genes identified as showing changes in translation, as compared in Figure S5A.
